# Turnover in life-strategies recapitulates marine microbial succession colonizing model particles

**DOI:** 10.1101/2021.11.05.466518

**Authors:** Alberto Pascual-García, Julia Schwartzman, Tim N. Enke, Arion Iffland-Stettner, Otto X. Cordero, Sebastian Bonhoeffer

## Abstract

Particulate organic matter (POM) in the ocean sustains diverse communities of bacteria that mediate the remineralization of organic complex matter. However, the variability of these particles and of the environmental conditions surrounding them present a challenge to the study of the ecological processes shaping particle-associated communities and their function. In this work, we utilise data from experiments in which coastal water communities were grown on synthetic particles to ask which are the most important ecological drivers of their assembly and associated traits. Combining 16S rRNA amplicon sequencing with shotgun metagenomics, together with an analysis of the full genomes of a subset of isolated strains, we were able to identify two-to-three distinct community classes, corresponding to early vs. late colonizers. We show that these classes are shaped by environmental selection (early colonizers) and facilitation (late colonizers), and find distinctive traits associated with each class. While early colonizers have a larger proportion of genes related to uptake of nutrients, motility and environmental sensing with few pathways enriched for metabolism, late colonizers devote a higher proportion of genes for metabolism, comprising a wide array of different pathways including metabolism of carbohydrates, amino acids and xenobiotics We find evidence in selected metabolic pathways for the existence of a trophic-chain topology connecting both classes. The interpretation of these traits suggests a distinction between early and late colonizers analogous to other classifications found in the literature, and we discuss connections with the classical distinction between r- and K-strategists.

## Introduction

The importance of understanding natural microbial communities in a scenario of global change is increasingly recognized [1]. In the open ocean, where external outputs of nutrients are scarce, it is estimated that 90% of the nutrients required to sustain primary production are obtained by the microbe-driven remineralization [2]. This collective activity of microbial communities can turn over particulate organic matter (POM) on the timescale of one week [2]. This rapid turnover makes it difficult to observe the processes occurring at the micro-scale. Since slight changes in the nutrients may have important consequences in processes such as CO2 sequestration, a better understanding of micro-scale processes will help to extrapolate predictions at a global scale [3].

Some of the difficulties in deriving general principles in the dynamics of POM are related to the high complexity of the nutrient composition of the particles [4] —which depends on the particle’s location, source, and size [5]. Also, *in situ* studies just very recently achieved single-day resolution [6]. For these reasons, while compositional and functional differences have been found when comparing lifestyles such as free-living vs. particle-attached bacteria [7, 8, 9], or locations, e.g. surface waters vs. deep waters [10, 11], little is known about the functions and micro-dynamics driving POM bacterial succession as particles are degraded.

To address these challenges, in previous work some of us studied the assembly of marine communities sampled from coastal waters on model particles, namely hydrogel beads composed of defined polysaccharides, or some combinations of these [12, 13]. Simplifying the heterogeneity associated with natural particles enables the study of highly diverse communities in a controlled manner.

In a first work [12], it was observed that community assembly in chitin particles happened in successions, and it was possible to differentiate three temporal phases: a short attachment phase (phase I: <12h), followed by a phase in which a drop in biodiversity was observed (phase II: 24h-48h), and a final phase in which biodiversity increased (phase III: 48h-140h). Metagenomic analysis of the assembly in chitin showed that bacteria in phase II had a higher frequency of extracellular chitinases, chemotaxis genes. In addition, some isolated strains were motile in laboratory assays.

In a second work [13], the number of substrates used were extended to alginate, agarose, carrageenan and some pairwise combinations of those. Similar succession patterns were observed to those found in the first work in chitin. Considering several resources allowed to observe that taxa in phase II were substrate-specific, while phase III was characterized by nonspecific taxa. This result suggested that the underlying environmental conditions determined the ecological succession to a great extent, and hence that selection was the dominant force, in particular in phase II were substrate-specific taxa were observed. Finally, in both works it was shown that isolated strains observed in phase III weren’t able to grow in the particle’s polysaccharide, but they were able to grow in the spent media of strains isolated from phase II, further identifying some of the metabolites consumed by late colonizers [13]. This suggested that facilitation was the main mechanism promoting the succession between phase II and phase III.

In this work we pursue two objectives. In the first place, we want to quantitatively reevaluate the interpretation associated with the observed phases (i.e. selection in phase II and facilitation in phase III), providing said interpretations with greater statistical support and encompassing them in a broader conceptual framework. For instance, it will allow us to discriminate the spatio-temporal scale in which selection operates, differentiating between selection at the metacommunity level, at the local community level, or at both levels. As a final corollary, the analysis will allow us to establish more direct connections between these experiments and observations from samples collected from the natural environment, which is often limited to looking for statistical patterns at the community level. This last point is of great importance for establishing connections between top-down patterns and bottom-up mechanisms established in controlled experiments.

To carry out this first objective, we re-analyzed the 16S rRNA sequencing datasets presented in Ref. [13], and incorporated new experiments on two additional substrates (chitosan and a combination of alginate and chitosan). We then followed a pipeline with well-established methods which combines neutral models [14, 15], with beta-diversity analysis and phylogenetic information [16]. Our results show that selection drives the assembly of the communities at both metacommunity and local community levels, and provide statistical support for the role of facilitation, further suggesting a direct interaction between primary degraders and secondary consumers.

Our second objective aims to explore whether the traits observed by analyzing metagenomic data in chitin [12] are also found in other substrates, given that the taxonomic specificity found in phase II would lead us to expect that there are specific traits for each substrate. To achieve this objective, we considered three new data sources: experimental metagenomics in all substrates presented in Ref. [13] and in the two new substrates incorporated here, metagenomic predictions with PICRUSt [17], and a dataset of 65 genomes sequenced from strains isolated in the two previous works [12, 13].

Similar to the analysis performed in Ref. [12] only for chitin we asked if, within the 8 substrates considered, we could find significantly different traits between temporal phases and, if they existed, if these were different to those previously found for chitin. We found that the most significant differences appeared when we compared phase I, phase II and phase III colonizers irrespective of the substrate in which they were sampled. Moreover, the phylogenetic autocorrelation decay was consistent with the temporal ranges in which the phases were defined, supporting that these traits may reflect specific adaptations.

Analysis of these traits showed that bacteria found in the selection phase consistently had a higher proportion of genes related to motility, chemotaxis and transporters. These traits, along with others such as genes related to ribosome biogenesis, suggested that these bacteria are specialized in the efficient search and uptake of immediately available nutrients and in rapid growth. On the other hand, the colonizers found in the facilitation phase have a higher proportion of genes involved in a wide variety of metabolic processes, including autonomous amino acid biosynthesis, carbohydrate metabolism or xenobiotics metabolism. Importantly, the differences that we find between the phases consider all the substrates aggregated, and are more and more significant than those that we find when comparing substrates. This suggests that there are functional groups associated with each stage of ecological succession, regardless of the substrate considered.

The distinction between both types of communities has connections with the distinction between rand K-strategists [18, 19]. The terms r and K refer to population dynamics parameters describing the maximum rate of increase and maximum equilibrium density of the population, respectively, r-strategists would dominate variable environments with abundant resources in which there is little competition, while K-strategists are adapted to environments with constant and scarce resources, where competition is harsh. This distinction has been widely exploited (and criticized) in macroscopic ecology [20]. However, its relevance in microbial ecology and, more specifically, the identification of r/K traits and their associated environments which would provide the grounds for a mechanistic justification, remain largely unknown [21]. Our work suggests that, in these experiments, ecological succession resembles a transition between rand K-strategists, which could be a general result for environments with intermittent inputs of resources.

## Results

### Illustrating microbial assembly signatures of ecological succession

Using synthetic model particles we studied community composition for both particles-attached and free-living bacteria in cultures incubated with seawater. Since part of the data used was previously published in Ref. [13], in this section we recover part of the analysis presented there for completeness, which will also helps us to illustrate the experiments.

The analysis of community composition revealed that the most abundant phyla were Proteobacteria (most notably *γ*- and *α*-Proteobacteria) and Bacteroidetes (mainly Flavobacteria). Microbes belonging to the *Halomonas* and *Shewanella* genera dominated the community in the earliest samples (labelled 0h) independently of the substrate used, suggesting that they encoded specific traits required for early colonization (Fig. 1). In the figures, the three replicates appear aggregated. Detailed trajectories for three substrates are presented in Suppl. Fig. 1 and interactive visualizations for all experiments are available in Suppl. Materials.

**Figure 1:**
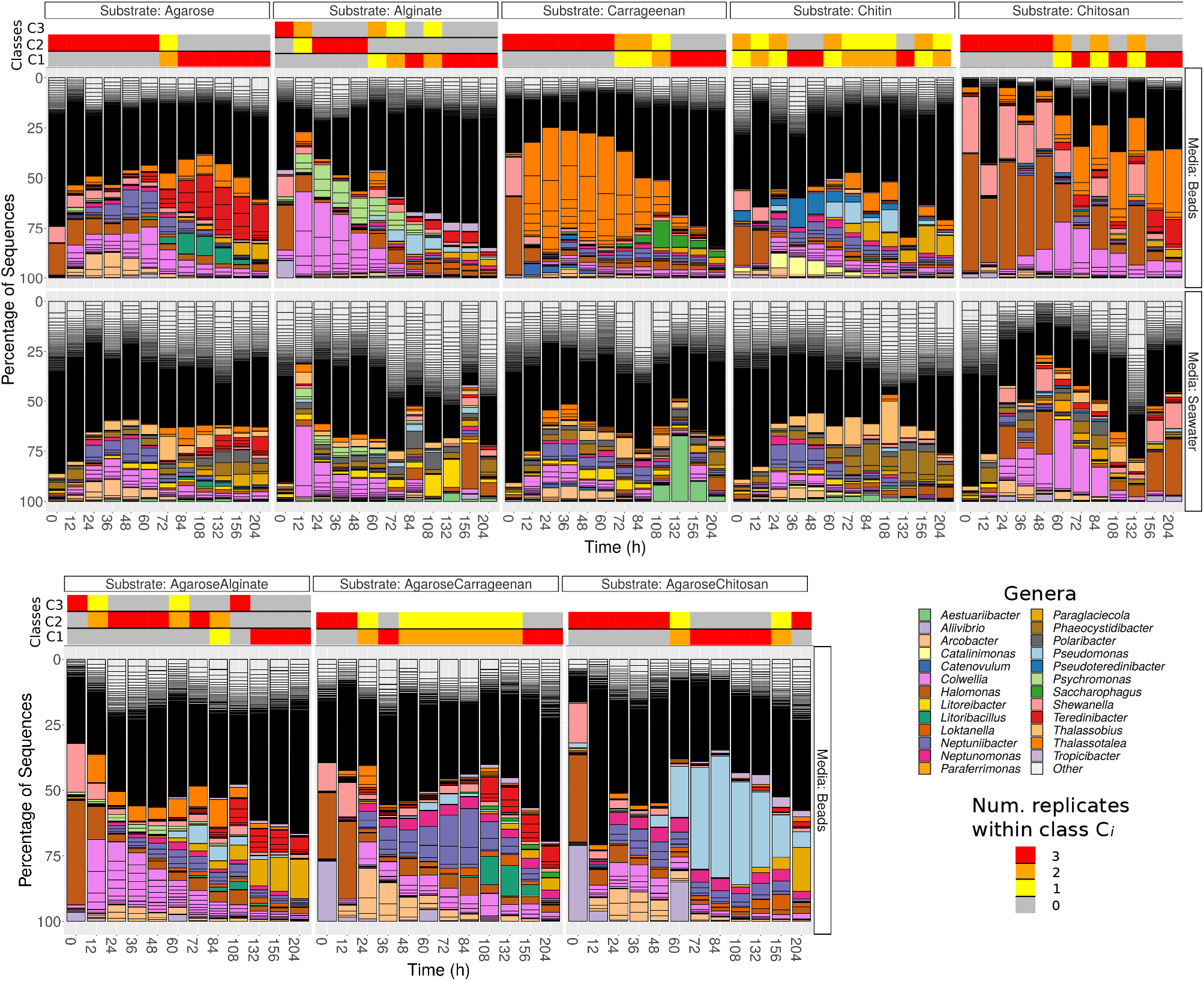
Relative abundances of genera for populations attached to beads made of pure substrates (first row) and present in surrounding water (second row) for each substrate (columns) at the different time points. The third row represents the population dynamics of beads made of mixed substrates. In the beads experiments, the horizontal bars on the top of the bar plots represent the classes obtained with unsupervised clustering, and the colour the number of replicates assigned to the different classes at each time point.

After 12h to 48h, the substrate-specific invasion presented in Ref. [13] is apparent from genera such as *Colwellia* and *Psychromonas* (alginate), *Arcobacter* and *Neptuniibacter* (agarose), *Thalassotalea* (carrageenan) or *Catalinimonas* and *Pseudoteredinibacter* (chitin). In chitosan, one of the new substrates incorporated in this study, *Halomonas* and *Shewanella* were slowly replaced by *Colwellia* and *Thalassotalea*. (Suppl. Fig. 2), which could have been caused by the fact that this substrate is more recalcitrant to decomposition. Another new pattern not reported in previous work is that bacteria found on the particles in high proportions were not highly represented in seawater before particle colonization (second row Fig. 1 and Suppl. Fig. 3 for the number of reads in seawater), and only increased in seawater after their proportion increased on the particles.

Although substrate-specific taxa – which in previous work have been shown to act as degraders [12, 13]–remained in the community at late time points (e.g. > 200h), there was a systematic increase in the diversity of the community driven by the invasion of substrate-unspecific taxa (see Suppl. Fig. 4), which were possibly unable to degrade the particles, but were successful in competing for metabolic byproducts. Interestingly, the trajectories of particle taxonomic composition over time projected onto the components of a principal coordinates analysis showed that these trajectories were similar irrespective of the substrate, and were driven by the temporal phase of succession (Suppl. Fig. 5), including the previous considerations for chitosan. These patterns suggest that resources determined a shift between substrate-specialist taxa dominating at early phases, when there was mostly a single homogeneous resource, and secondary consumers colonizing the particles when the number of micro-niches increased.

### Unsupervised classification finds classes aligned with the temporal phases

The above patterns suggested that selection dominated the global assembly of these communities. Nevertheless, it may be possible that selection is only operating at a metacommunity-level with the local assembly being stochastic. To test this hypothesis, we developed an approach combining well-established methods which determined whether the assembly of the communities was best explained by stochastic or selective processes and, under the latter scenario, determined which were the most important selective forces. The approach and hypothesis are summarized in Table 1.

**Table 1:**
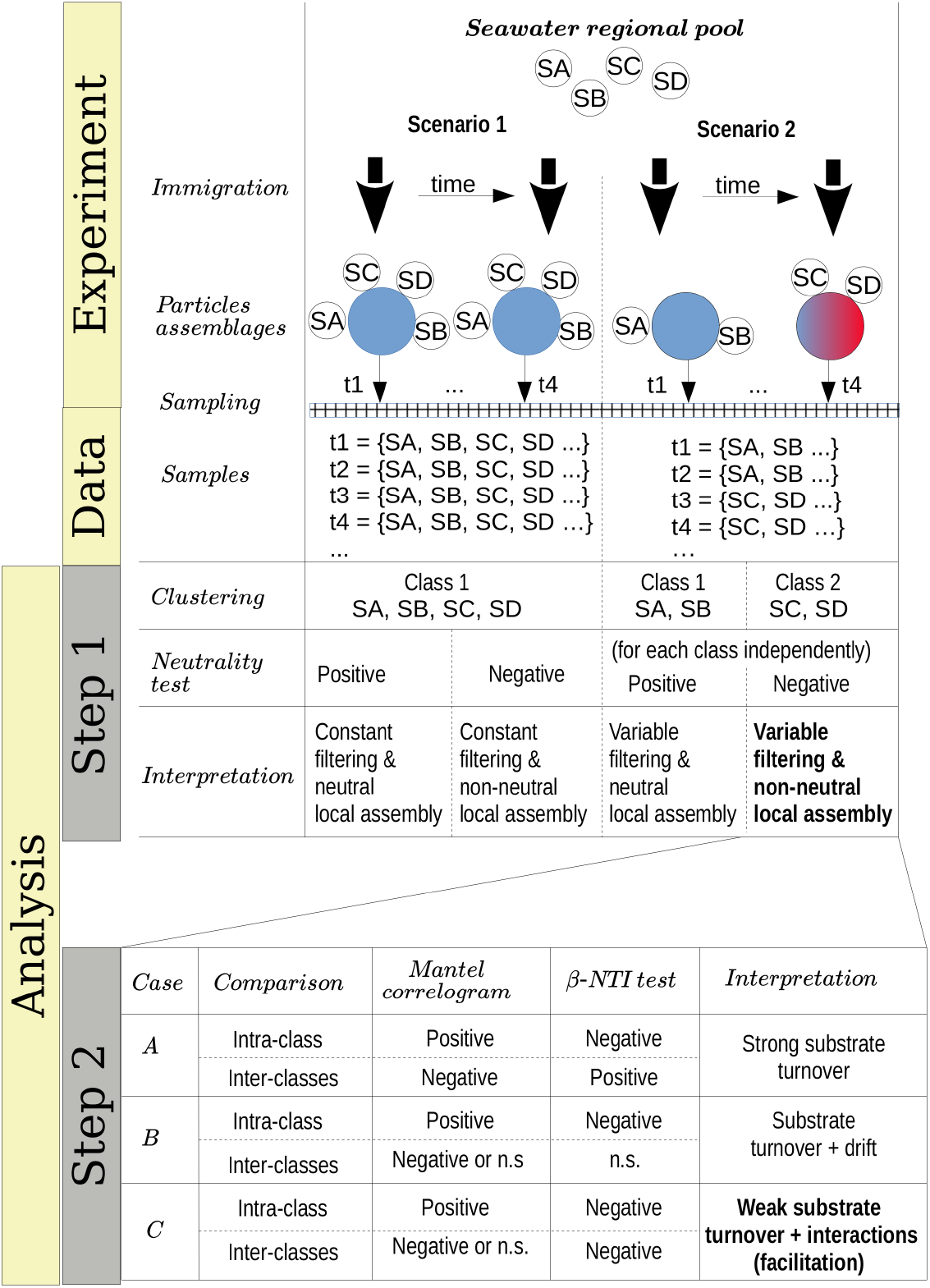
Methods and hypothesis. Our approach proceeded in two steps. (Step 1) For each substrate, immigration from the seawater species (circles labelled as “SA”, etc) onto the beads (coloured circles) could be stochastic or driven by selection. We started classifying the beads’ samples on the basis of their *β*–diversity similarity. If we did not find clusters (scenario 1) we performed a test of neutrality considering all the samples of that substrate. If we found clusters “community-classes” (scenario 2), we investigated if it was a signature of distinct selection forces acting in different classes (“metacommunity selection”). Under this scenario, neutrality was tested considering each subset of samples determining each class independently, hence creating a different metacommunity for each of them. The interpretation depending on the different outcomes is indicated, with the results we found highlighted in bold in the table. (Step 2) Since selection at the metacommunity and local levels was suggested, we investigated its mechanisms considering the phylogenetic similarity of the samples. The combination of results from Mantel correlograms and *β*–NTI comparisons led to three cases compatible with our results in Step 1. The analysis of these cases suggested that the dominant processes were those indicated in case C and, in a lesser extent, case B. Very few comparisons supported case A. Note that these cases were restricted to those compatible with our results in Step 1 and other possibilities (e.g. homogenizing dispersal) were excluded.

Unsupervised classification of the communities according to their compositional similarity (see Methods and Suppl. Fig. 6) revealed that the optimal classification divided the communities into two classes, with the exception of the experiments on alginate and on a mix of agarose and alginate, which resulted in three community-classes. The classification found for each substrate is indicated in the coloured horizontal bars on the top of the bar-plots in Fig. 1. The classes found could indicate the presence of selective forces acting similarly on subsets of samples. Consistent with previous work [12, 13], the classes matched temporal phases: the method systematically clustered all three replicates of samples collected at early time points (phase II: 12-60h) in one class, and samples from late time points (phase III: 108-204h) in a another class. Samples at intermediate time points (dependent on the substrate, between 60-108h) were sometimes split between each class. Hereafter, we term the interval 60-108h “transition phase”. The additional class found on alginate and the mix of alginate and agarose corresponded to early samples (phase I: 0-12h, “attachment phase’’) and to the transition phase. As an exception, we found that chitin classes were not sharply separating both phases, a result that would not be consistent with previous work [12] but that we attribute to experimental noise. We observed that samples in the first class of chitin had a significantly lower number of reads than the rest of the particles’ dataset: median 4838 (IQR: 3191-8479) vs. 175605 (IQR: 66421-326622). Results presented below support this explanation.

### Selection dominates at both metacommunity and local-community levels

The method we used to cluster the communities ([15]) was framed in the context of neutral theory, with each class being interpreted as a set of local communities whose species members immigrate from the same metacommunity (see Methods and Table 1, Step 1). Since the community classes are compositionally differentiated, in the context of Hubbc?s model of neutral assembly we shall expect that a different speciation rate occurs in each class (interpreted as a metacommunity). This is a conservative approach since rejecting neutrality will be more difficult considering a separate parameterization for each metacommunity –in other words, a single metacommunity will be unable to generate local samples belonging to the different configurations determined by the different classcss [14]. To test if these classes were compatible with a neutral assembly we fitted, for each class within each substrate, a model to a Hierarchical Dirichlet Process (HDP) with the software provided in Ref. [14]. Following this method, the likelihood of the observed communities of being neutral was compared with the likelihoods of artificial communities generated with the fitted HDP (see Methods). We tested two possibilities, one in which both the metacommunity and the local communities are assembled through a neutral process (complete model) and another one in which the local communities but not the metacommunities are assembled neutrally (local model).

15 out of 18 classes rejected the neutral hypothesis (empirical-*p* < 4 × 10^-4^), supporting the observation that the species were not functionally equivalent within each class and that both the metacommunity and the local community assembly are driven by selection (see Table in Fig. 2). Predicted ‘neutral’ community classes included the third community class identified on alginate particles, which colonized these particles from time 0-12h (empirical-*p* > 0.20); the first community class on chitosan (0-48h, marginally significant; empirical-*p* = 6 × 10^−4^ for the local model); and chitin was again an exception, with the first community class appearing neutral (defined along the whole trajectory, empirical-*p* > 0.8). We believe this abnormally high empirical-p value for the first class on chitin reflects the low copy number found for this class, which suggests it may be an artefact. The other two remaining examples (first class on chitosan and third class on alginate) suggested that species may only be considered functionally equivalent at very early phases of the assembly (“attachment phase”), with chitosan having a longer attachment phase.

**Figure 2:**
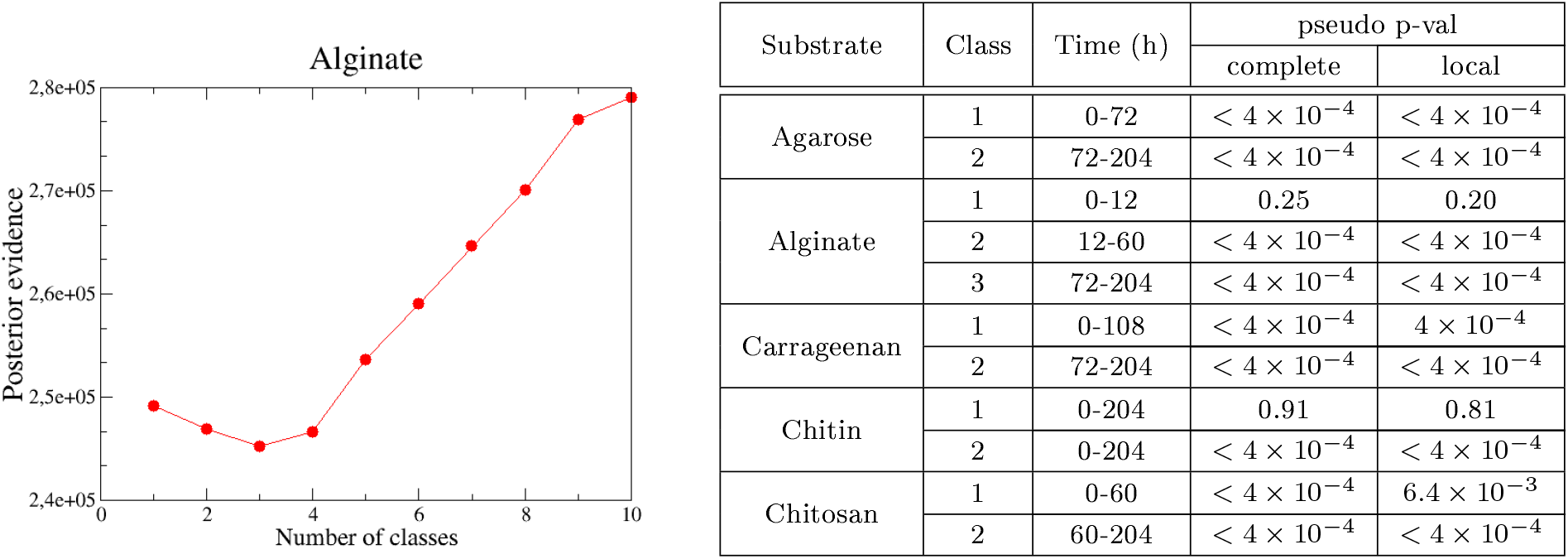
Number of optimal classes and test of neutrality. (Left) Optimal number of classes (number of Dirichlet mixture components) are determined identifying the minimum of the posterior evidence of the fit. Results are for alginate. (Right) Main time intervals determined by each class in single substrates. The time window is determined considering consecutive samples classified in the same class (i.e. sparse samples misclassified are non indicated, see Fig. 1 for details). For chitin the whole interval is indicated since no clear time-windows are defined. Pseudo p-valucs of the HDP test for each class found in pure substrates for the complete and local models are indicated.

Beyond these exceptions, we found 15 classes with a good correspondence with phase II (12-60h) and phase III (108-204h) which rejected neutrality, suggesting that they were mainly driven by selection. These results provide a solid support for the observations made in previous work, further showing that selection operates at both metacommunity and local levels. We also show that selection not only operates at phase II but also at the more taxon-unspecific phase III.

Since there was a fair overall agreement on the determination of the classes across substrates and on their correspondence with time phases, to simplify the following analysis we directly focused on differences between phases: phase I (0-12h), phase II (12-60h), and phase III (108-204h). We excluded from the analysis the transition phase (60-108h).

### The succession is not driven by a complete substrate turnover

After determining that selective processes have the most important role in assembly in most experiments, we further investigated the determinants of selection (see Table 1, Step 2).

We first estimated how strong is the phylogenetic correlation between communities and which is their characteristic temporal decay. In particular, we would like to know if this decay is consistent with the temporal phases identified with unsupervised clustering, which would provide us with firm support to interpret traits associated with these phases as a phylogenetic signal. We estimated the phylogenetic distance with Unifrac, which is a *β*–diversity metric incorporating phylogenetic relatedness [22]. We computed the correlation between Unifrac distance and temporal distance that separated each pair of communities by computing Mantel tests. To estimate the correlation decay, we binned the comparisons in subsets of increasing temporal distances, performing an independent Mantel test for each subset (i.e. a Mantel correlogram [23], see Methods).

As expected, we found that communities closer in time were phylogenetically more similar (see Fig. 3A for alginate and Suppl. Fig 7 for other substrates) and that, when the temporal distance between samples increased, the Mantel statistic became non-significant (Fig.3A) or even significantly negative on some substrates (Suppl. Fig. 7). Importantly, the results show that the phylogenetic similarity is significant within the same phase as soon as the communities are closer than 40h in time. This give us support to interpret traits within the same phase as a signature of adaptation. Notably, the minimum temporal distance between phases II and III (i.e. the width of the transition phase) is 48h, a distance within which the phylogenetic signal decayed, indicating that there was significant phylogenetic turnover between both phases.

**Figure 3:**
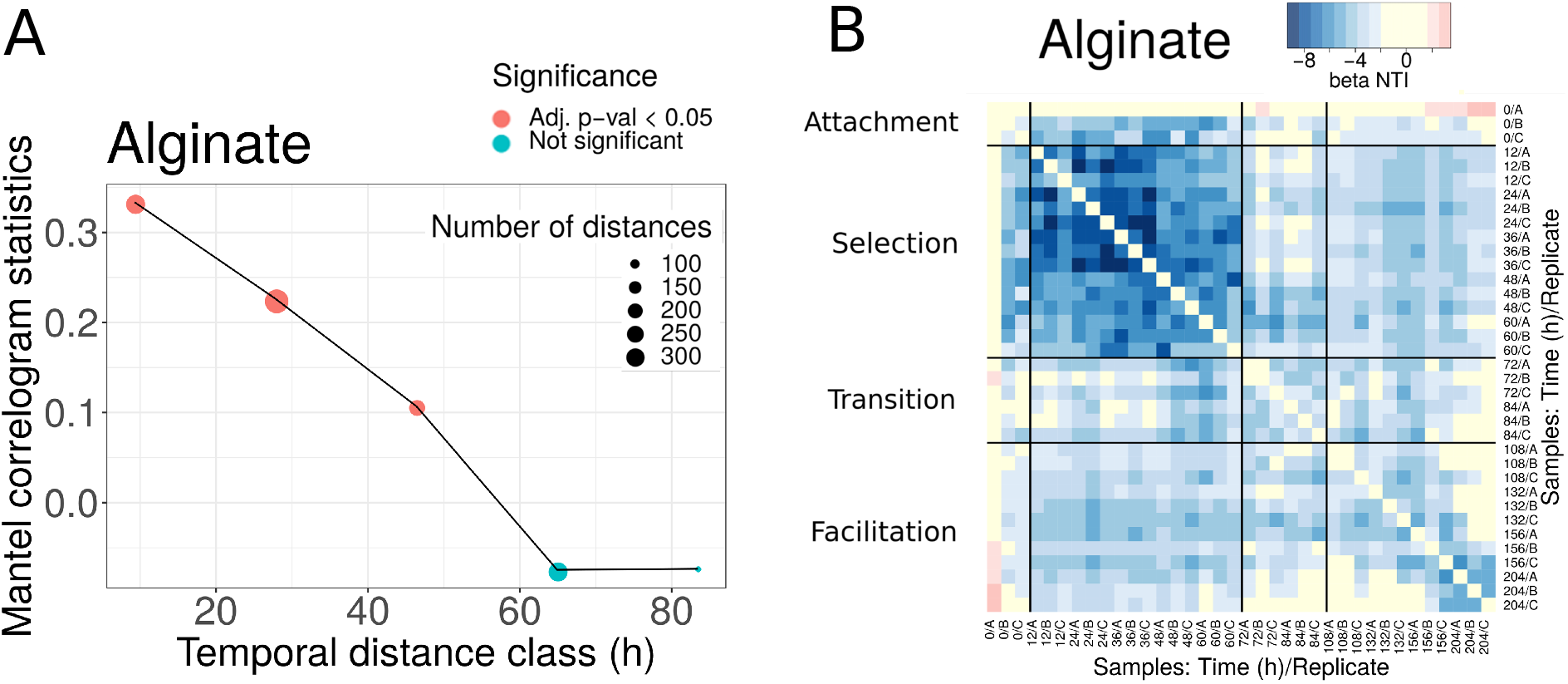
Phylogenetic turnover for experiments in alginate. **(A)** Mantel statistics indicating the correlation between the phylogenetic and time distances in beads of alginate. Both matrices are split in subsets corresponding to different ranges of distances, and an independent test performed for each subset. The middle point of each range is indicated in the x-axis, and the number of distances in each subset and the significance of each test is shown in the legend. (B) All-against-all comparison of the *β*NTI index (excluding diagonal values). The black thicker lines separate comparisons within and between the different phases. Significant values correspond to |*β*NTI|>2, non-significant values are shown in yellow. Results for other substrates are provided in the Suppl. Material.

The second question that we would like to study through the analysis of statistical phylogenetic patterns is whether we can evaluate the relative importance of resource transformation, direct interactions, and drift, in ecological succession. We consider two extreme scenarios. In the first scenario, the degradation of the main substrate could lead to a complete substrate turnover. Under this scenario we would expect a complete phylogenetic turnover, indicating that the communities present in the selection phase abandon the particles before the colonization of the communities present in the facilitation phase occurs. Therefore, the two types of communities would have little coexistence, in which case any facilitation would be indirect. In the second scenario, this transformation of the environment is not so strong and the communities in the facilitation phase would be assembled in direct interaction with members of the communities in the selection phase, which remain through the trajectory. The Mantel correlogram decay observed and the compositional turnover would *a priori* favour the first scenario.

To investigate these questions, we computed the *β*–Mean Nearest Taxon Distance (*β*MNTD) [16]. For each pair of communities A and B, the *β*MNTD was computed by measuring the phylogenetic distance of each taxon in A with respect to its closest relative in B, and then by averaging across all taxa (and symmetrically for B). Then the taxonomy was shuffled a large number of times to estimate the null expectation and a z-score computed, termed *β*-Nearest Taxon Index (*β*NTI) [16] (see Methods). A *β*MNTD value significantly higher than the null model (*β*NTI > 2) is expected when the community turnover is driven by a strong shift in the environmental conditions (termed variable selection [24]). On the other hand, a significantly negative value (*β*NTI< −2) indicates that strong and homogeneous environmental conditions make the communities taxonomically more similar than expected by chance (termed homogeneous selection [24]). Therefore, this metric may allow us to narrow down the hypothetical scenarios described above, by comparing the results we found for communities within the same phase and between different phases (summarized in Table 1, Step 2).

We found that the *β*NTI values were significantly negative (comparisons in blue in Fig. 3B for alginate and Suppl. Fig. 8 for other substrates), pointing towards a homogeneous environmental filtering as the main driver of the assembly. The strongest selection was observed at phase II (12-60h). This is an expected result, since we know that at early stages dominate taxa that are able to degrade the correspondent polysaccharide. Strikingly, in alginate (Fig. 3B) and agarose (Suppl. Fig. 8) this signal was also observed in the comparisons between phase II and phase III communities (comparisons within distant off-diagonal boxes). For other substrates such as chitin or carrageenan (Suppl. Fig. 8) the comparison between phases revealed that drift had a major role in the transition, but there was no evidence of complete turnover of resources either.

These results were in apparent contradiction with the compositional turnover and the Mantel correlogram decay. However, since the *β*NTI focuses only on the closest relatives, these results are explained if there were taxa from phase II remaining in phase III (if a given taxon was present in both early and late communities its contribution to the *β*MNTD was zero). The explanation compatible with both a significant compositional turnover and the permanence of some taxa along the assembly was that members of communities in phase II and communities in phase III coexisted, and that the new microniches needed for the compositional turnover were continuously generated by resources produced by those degraders remaining from phase II. This indicates that facilitation occurs in direct interaction, as opposed to an indirect process in which the resources were fully transformed in phase II and then consumed by new colonizers in phase III. Aligning our analysis with results from previous work, in the following we will denominate phase II “selection phase” and phase III “facilitation phase”.

### Metagenomic analysis reveals differentiated ecological strategies throughout the succession

Our previous results suggested that there was phylogenetic turnover, driven by a transition between strong environmental selection at early time points (selection phase) and a combination of environmental selection and ecological interactions at late time points (facilitation phase), with marginal evidence of drift. We aimed to investigate if this distinction translated into signatures in the genetic repertoires of the communities at the different phases, hence reflecting specific adaptations. For this study, one sample per time point was collected in one of the replicates and its metagenome sequenced.

The number of reads in the metagenomics data increased with time (Suppl. Fig. 9). However, the number of reads was generally lower than for 16S sequencing data, and we discarded samples with too few reads, mostly affecting the attachment phase (see Methods). To complement this information, we performed a prediction from the 16S rRNA amplicon sequences with PICRUSt [17], from which three replicates per sample and time-point were available. We provide in Suppl. Materials an evaluation of the accuracy of the PICRUSt predictions. In the following, we focus on those results consistent between metagenomic data and the metagenomes predicted from 16S rRNA with PICRUSt. Analysis of the attachment phase was performed only from the predictions.

A principal component analysis of the metagenomic predictions revealed that the attachment, selection and facilitation phases were projected in orthogonal directions, suggesting the existence of distinctive traits in each phase (Fig. 4A). On the other hand, when the same data representation was considered and the samples were coloured according to the substrates on which the communities assembled, no clustering was apparent (Suppl. Fig. 10). Indeed, comparing the difference in the mean proportion of genes belonging to different substrates, few differences were found and most of them were against chitosan which, as we showed, seemed to have some delay in the colonization (see e.g. comparison against alginate in Suppl. Fig. 10). Hence, to investigate the existence of distinctive traits, we aggregated all substrates and compared the difference in the mean proportion of genes belonging to samples in the selection phase against those in the facilitation phases for both the metagenomics data and the predictions (Fig. 4B and C).

**Figure 4:**
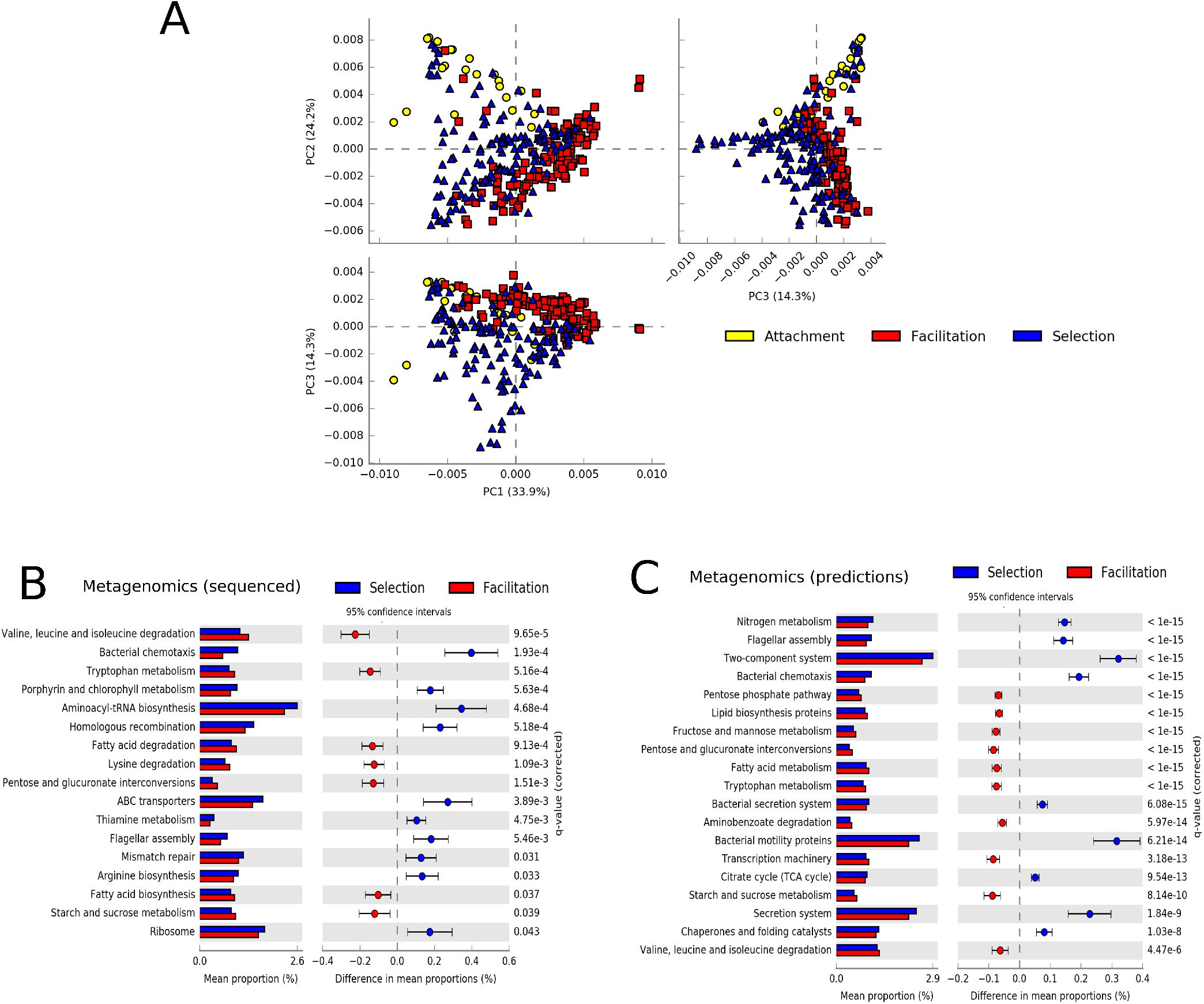
Summary of metagenomic features. (A) Principal component analysis of the predicted metagenomic profiles coloured by the phase in which they were sampled. (B and C) Difference in the mean proportions of genes between communities at the selection and facilitation phases in metagenomic experiments (B) and in PICRUSt predictions (C). Each row in the diagram represents genes classified in the KEGG pathway indicated. The first column represents the mean proportions of the genes in the pathway for each phase, and the second column the difference between those proportions. Adjusted Benjamini-Hochberg p-values and 95% CI intervals are indicated. Pathways with effect sizes lower than 0.1 (B) and 0.05 (C) were filtered to show a similar number of pathways.

Both experimental metagenomics and PICRUSt predictions provided a similar qualitative picture. Colonizers at the selection phase predominantly encoded pathways related to motility and environmental sensing (e.g. bacterial motility, two-component systems), replication and repair (e.g. homologous recombination, mismatch repair), translation (e.g. aminoacyl-tRNA biosynthesis, ribosomal genes –which are characteristic of fast-growing bacteria [25, 26]) or transport (ABC transporters, bacterial secretion systems). Colonizers at the facilitation phase showed more genes related to metabolic pathways. More specifically, genes related to carbohydrate, lipid and amino-acid metabolism were more represented in the facilitation phase. There were, however, a few metabolic pathways more represented in the selection phase such as the metabolism of vitamins and cofactors, arginine biosynthesis, and the metabolism of nitrogen. Notably, some pathways were consistently predicted from both 16S rRNA and metagenomic datasets for both early colonizers (e.g. flagellar assembly, bacterial chemotaxis) and late colonizers (e.g. pentose and glucuronate interconversions, valine, leucine and isoleucine degradation). Importantly, there were no pathways yielding contradictory information (i.e. predicted as enriched in one successional phase in one dataset and in another phase in the other dataset).

We finally compared the difference in mean proportion of genes between samples in the attachment and facilitation phases for the metagenomics predictions only (Suppl. Fig. 11). Although some of the pathways enriched in the attachment phase were similar to those found at the selection phase (e.g. bacterial chemotaxis, flagellar motility) the most characteristic feature of the attachmentis phase was the large proportion of genes related to transporters, in particular ABC transporters, which ranged from 0.2% for samples at the selection phase in the experimental metagenomes to a 0.6% in the attachment phase for the predictions.

### Ecological strategies are imprinted in the genomes of isolated bacteria

To test previous results and to gather insights into the relation between taxa and traits associated with each community phase, we analysed annotated draft genomes of 65 isolates derived from particle communities [12, 13]. We combined 16 assemblies from previous work with additional 49 particle-derived genomes to arrive at a set of genomes representing major ESVs present in particle-associated communities. We assigned the isolates into classes depending on the phase of colonization in which they had a high propensity for being observed. Isolates’ propensity was determined from the propensity of the ESV having a 100% sequence identity with the 16S isolate gene. Propensity of the ESVs were estimated analysing their relative abundance in the assembly experiments (see Methods). More specifically, for each ESV we computed a generalized linear model with phases (attachment, selection and facilitation) as predictor variables and their abundance as a response (see Methods). Significant coefficients indicated propensities for specific phases, and these propensities were assigned to the corresponding isolate to determine the classes (see Suppl. Fig. 12). We subsequently compared the functional gene content of genomes in each class.

All isolates were classified as having either a preference for the attachment (6), selection (10) or facilitation phases (13), with the remainder being classified as generalists (i.e. a preference for at least two phases, in most cases including selection and facilitation phases). Only two isolates were predicted to have no significant assignment (see Fig. 5A). Isolates classified in the attachment phase belong to the *Vibrionaceae* family, and most of those classified at the selection phase belong to *Alteromonadaceae*. In contrast, the facilitation phase has members of several families, most of them belonging to the *Flavobacteriaceae* and *Rhodobacteraceae*. In the phylogenetic tree (Fig. 5A) it was apparent that generalists species were phylogenetically closer to those isolates classified in the facilitation phase, suggesting that their ecological strategies could be more similar.

**Figure 5:**
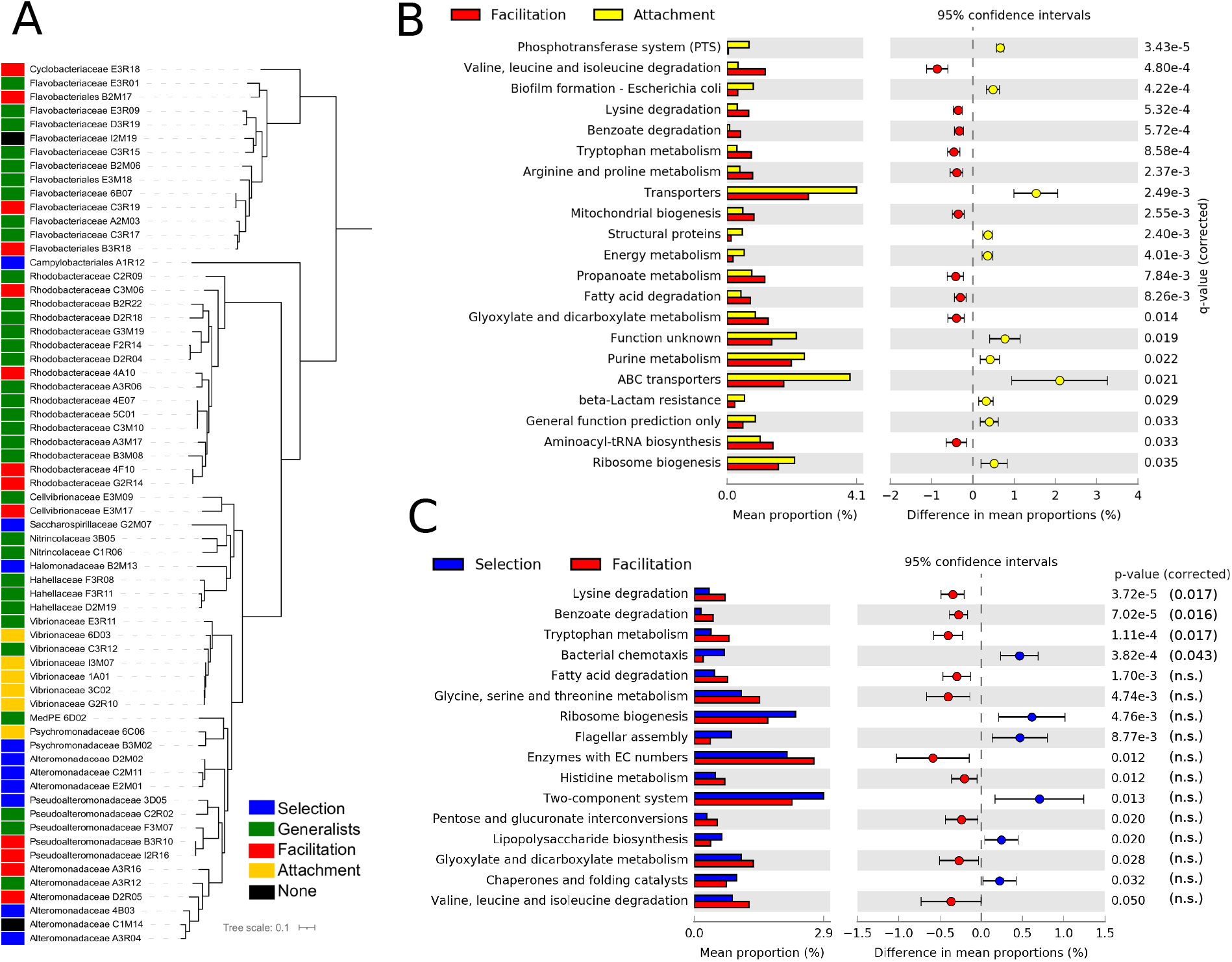
Ecological strategies of isolated strains. **(A)** Phylogenetic tree of the isolates used in this study. The ecological strategies of the isolates inferred from the trajectories of the ESVs with 100% sequence identity in the 16S amplicon sequencing experiments are shown with different colors. (B) and (C): Differences in the mean proportions of genes grouped in KEGG pathways between isolates identified as having a preference for the attachment phase and those with a preference for the facilitation phase (B) and the same comparison between isolates with a preference for the selection phase and those with a preference for the facilitation phase (C). The first column represents the mean proportions of the genes in the pathway for each group of isolates, and the second column the difference between those proportions. Adjusted Benjamini-Hochberg p-values and 95% CI intervals are indicated. Only pathways with effect sizes larger than 0.1 are shown.

The gene content comparison between isolates classified in the facilitation phase and in the attachment phase yielded the most significant signal (Fig. 5B). Consistent with the metagenomics analysis, we observed that a typical facilitation genome tended to encode a higher proportion of central metabolic pathways related to degradation, such as pathways to break down amino acids and fatty acids. Also consistent with metagenomic predictions (Suppl. Fig. 11), the most notable feature of attachment genomes was the high proportion of transporters, in particular ABC and PTS transporters (Fig. 5B). Other pathways that were significant in the metagenomes, such as those related with genetic information processing (e.g. chaperones and folding catalysts), were also encoded by a higher proportion of attachment genomes (Fig. 5B).

Focusing on the comparison between the selection and facilitation phases (Fig. 5C) we retrieved a similar picture to the one found in the metagenomics data. However, the significance was lower, and most of the pathways were not significant when the p-value was corrected for multiple testing, possibly due to the low number of isolates considered in both groups.

Taking together both metagenomes and isolates, we found a consistent distinction in the genetic signatures characteristic of the different successional phases, summarized in Suppl. Tables 1, 2 and 3.

### Analysis of specific pathways reveal a trophic-chain topology

We explored in more detail some specific pathways to gain a finer scale understanding of the core traits characteristic of the different community types. When narrowing down to specific genes, we focused on the set represented in a significantly higher proportion in metagenomic data from one of the successional phases. Since the metagenome-associated signal may be dominated by the contribution of few very abundant species, we quantified the proportion of isolates from each class that encoded each gene in a pathway to assess the extent to which traits were shared broadly by taxa in a class. Results are summarized in Fig. 6.

**Figure 6:**
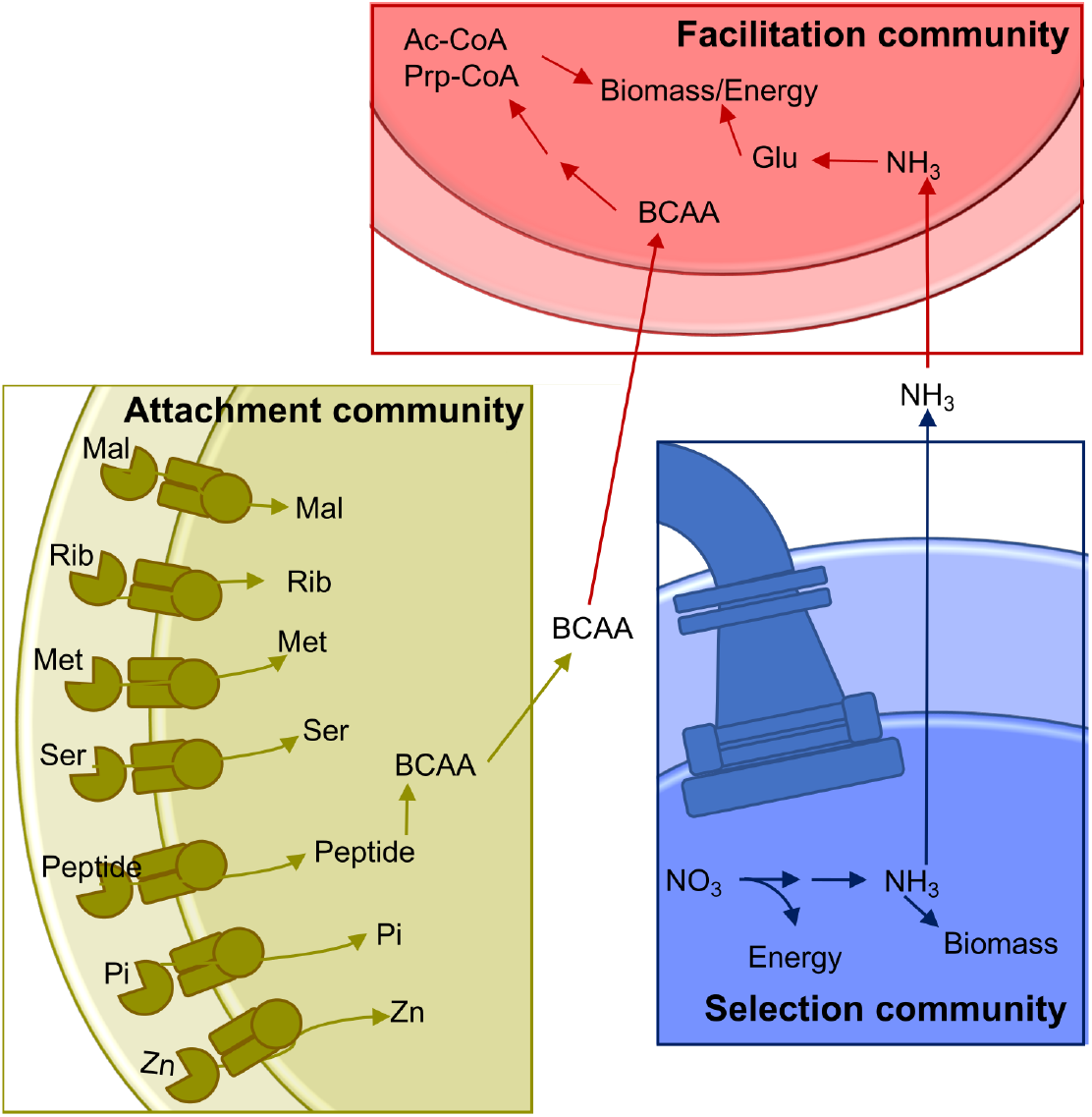
Analysis of specific pathways. Schematic of enzymatic processes encoded by taxa belonging to the three community types that assemble on polysaccharide particles. Attachment communities (gold), that are the first to form on particles and commonly encode a diverse array of high-affinity ABC transporters, specific for sugars such as maltose (Mal) and ribose (Rib), amino acids including methionine (Met) and serine (Ser), transporters specific to small peptide signals/antimicrobials, inorganic phosphate (Pi), and zinc (Zn). They also have genes to synthesize BCAA. Selection communities (Blue) form in between attachment and facilitation communities. Taxa in these communities frequently encode flagella and chemotaxis genes, and perform dissimilatory nitrate reduction generating ammonia. Facilitation communities (Red) are the last to form on particles and encode pathways that break down BCAA into acetyl-CoA (Ac-CoA) and propanoyl-CoA (Prp-CoA), metabolites that can enter central metabolism to make biomass or energy. Taxa in facilitation communities also commonly encode glutamine and glutamate (Gln) synthases and glutamine synthetase, which are the main pathways for assimilation of ammonia-derived nitrogen into biomass.

Attachment communities encoded a high proportion of ABC-transport associated genes relative to other community types (Figs. 4 and 5). We found genes encoding 23 ABC transport complexes represented in higher proportion in the attachment-associated metagenomes (yellow boxes in Suppl. Fig. 13); 16 of the predicted complexes had at least one gene more represented in the isolates associated to attachment than in any other isolates class, suggesting that these ABC transporters are commonly encoded by isolates in the attachment communities (shown with asterisks in Suppl. Fig. 13). Among these complexes, we find transporters predicted in a variety of bacteria to encode high-affinity uptake systems such as transporters for maltose [27], ribose [28], methionine [27], serine (Aap-JQMP genes [29]), inorganic phosphate [30], zinc [31], and peptides that can act as signals or antimicrobials (Yej-ABEF genes [32])(Fig. 6).

Interestingly, we also found ABC transporters for branched-chain amino acids (BCAA) that were significantly enriched in the facilitation community metagenomes (red boxes in Suppl. Fig. 13), and in isolates genomes (asterisks). Following up on this finding, we found that, in addition to encoding BCAA transporters, facilitation phase genomes encoded a higher proportion of the genes for valine, leucine and isoleucine degradation than isolates classified in different community phases (asterisks in Suppl. Fig. 14), a finding that agreed with metagenome predictions (Suppl. Table 3) and PICRUSt predictions (red boxes in Suppl. Fig. 14). BCAA degradation produces metabolites that feed into central metabolic pathways for energy generation and biomass. The ability to break down valine, leucine and isoleucine to acetyl-CoA, succinyl-CoA and propanoyl-CoA, means that these amino acids can be used for the biosynthesis of lipids, sugars or other amino acids, and for energy generation through respiration[33](Fig. 6).

Having reinforced functional traits that characterized taxa in each community phase, we next looked for pathways that were split among phases, we hypothesized that such split pathways would link successional phases and represent evidence for trophic interactions between phases. In Suppl. Fig. 14 we observed that some genes with annotations related to BCAA overrepresented in the attachment phase (yellow boxes) formed a coherent module for the biosynthesis of valine, leucine and isoleucine (Suppl. Fig. 15). The overrepresentation of these biosynthetic genes in attachment communities, alongside the high proportion of BCAA transport and degradation genes in facilitation taxa is consistent with the idea that the availability of these amino-acids could facilitate the transition between successional phases (Fig. 6).

We also found evidence of pathways splitting in the nitrogen metabolism KEGG pathway. This pathway was the most statistically significant metagenomic prediction from PICRUSt, and was predicted to have a significantly higher mean proportion in selection communities than in facilitation communities (Fig. 4, first row). An analysis of nitrogen metabolism pathways in isolate genomes revealed clear differences in the ability of taxa from the three community types to metabolize different forms of inorganic nitrogen. Metagenomes and genomes from selection and attachment communities often encoded pathways of nitrate reduction (Suppl. Fig. 16), with the presence of these genes in the isolated genomes (indicated with asterisks) more consistent with dissimilatory nitrate reduction. In contrast, facilitation communities had higher proportions of genes to assimilate ammonia, a reduced form of nitrogen. We also observed high proportions of glutamine synthetase in facilitation community metagenomes, whose presence was confirmed in the genomes (node 6.3.1.2 in Suppl. Fig. 16) Glutamine synthase uses ammonia to catalyze the production of L-glutamine in an ATP-dependent reaction. The predicted production of ammonia from nitrate reductase activity in selection and attachment communities suggesting that the transition between selection and facilitation communities may be promoted by the conversion of nitrate to ammonia by taxa that assemble in selection communities (Fig. 6), adding another possible trophic link between community phases.

## Discussion

In this article, we applied a comprehensive computational pipeline on different types of sequencing data, aimed at understanding the relative role of stochastic vs. deterministic processes in microbial assembly, and at identifying the main traits involved in ecological succession. In previous work, we found that two well-defined phases were apparent in the ecological succession of these communities on particles [12, 13]. Despite the reproducibility of this transition, there were different scenarios in which deterministic and stochastic processes could operate explaining these patterns, and our pipeline proceeded step by step to narrow down the different hypotheses, following Ref. [34].

Firstly, we reanalysed 16S rRNA amplicon sequencing data, and we applied unsupervised clustering to detect compositionally similar communities [15]. The search for compositionally similar communities is a simple and powerful tool used in the past to identify human microbiome enterotypes [35], and to identify differentiated functional performances in experimental assays for tree-holes communities in beech trees (phylotelma) [36].

We found that, in all substrates, it was possible to identify two to three community-classes which, in most cases, clustered together communities sampled at early time points in one class and at the late phases of the colonization in the other one. The remaining class, when it was present, clustered samples in the very early phase or in the transition between early and late phases. This result suggested that the degradation of resources was driving the succession, possibly with an abrupt shift in the composition leading to the formation of distinct classes of communities. In other words, the community-classes would be shaped by the resources available at the particles, effectively exerting a filter leading to two differentiated classes.

Still, among those bacteria overcoming these filters, those within the same community-class could assemble stochastically, i.e. following the framework of neutral theory, members of the same community-class might be “functionally equivalent”. We tested this possibility by comparing the observed communities with communities generated with a neutral model with the method proposed in Ref. [14], finding that the abundances of the observed communities were significantly different from the expectation under neutral assembly. Therefore, selection was not just acting at a “metacommunity level” (in our experiments, within the time-frame in which each class is defined) but also at a “local level” (within each specific community).

There is considerable debate on the relative importance of niche and stochastic effects in natural bacterial communities, because it depends on a number of variables. In locations with harsh environmental conditions such as in the Antarctica, environmental factors dominate the assembly of microbial communities [37]. This is not always the case since, in the deserts, the observation was distinct for autotrophs (stochastic) and heterotrophs (selection) [38]. These patterns are also likely driven by the extent to which the environment is spatially structured. For instance, soil communities seem to be mostly structured by abiotic factors (in particular, pH) [39], with communities living in finer-grained sediments experiencing strong selection while, for shallow sediments more exposed to perturbations, stochastic processes are more important [16]. In the ocean, large-scale compositional patterns are related to environmental factors, such as temperature or salinity [40]. At the different depths, environmental filtering has been observed to be more important in shallow waters while dispersal limitation tend to be more important in deep waters [41]. Interestingly, another study considering natural samples and conducting a similar analysis to the one reported here showed evidence of selection at the metacommunity level in both surface waters and waters at 200m. [42], consistent with the existence of differentiated classes we also found. They also observed non-neutral assembly for the local communities in surface waters, consistent with our findings for the selection phase. But similar to the results presented in Ref. [41] they found support for neutral assembly for the local communities at 200m. If an analogy between the colonization occurring along the water column in the ocean and our experiments would be valid, the dominance of stochastic processes at late times would represent a difference with our results. These differences could be explained by the fact that our communities were sampled from coastal waters and assembled in controlled conditions on simplified particle substrates, possibly favouring selection processes. Indeed, the importance of random perturbations, which are frequent in the ocean, favour a stochastic assembly and can modify biogeographic patterns [43].

The reproducibility of the experiments allowed us to investigate further the mechanisms of selection. Under the conditions of our experiment, if there was a dramatic shift in the underlying resources, we would expect the compositional and phylogenetic similarity of the communities to diverge along the experiment, whereas communities at adjacent time-points should be more similar. We indeed found that the overall similarity was higher for adjacent time-points, and it was reduced for communities distant in time. Moreover, when we focused only on the likelihood of finding closest relatives between two communities with the *β*NTI metric [16], it was particularly high at early time-points (12-60h), suggesting that the environment is exerting a strong selection, making these communities more similar [24]. However, finding close relatives among distant communities was also more likely than expected by chance, even if there was significant phylogenetic turnover. This may be seen as an unexpected result, because the resources are degraded along the succession, and a complete turnover of the resources could be expected, as it would explain the compositional turnover. This is a scenario termed environmental variable selection that should lead to values of *β*NTI that are significantly positive [24]. Having discarded stochasticity, drift, and variable environmental selection as determinants of the shift, the Mantel correlogram and the significantly negative *β*NTI values are consistent with a scenario where initial resources have not been exhausted, allowing some degraders to stay on the particles along the path, and these provide new resources for the colonization of new species on top of them. The results are therefore aligned with the mechanistic experiments presented in previous work showing that facilitation allowed to some isolated strains to grow on media in which the polysaccharide used in the experiment was the sole carbon source [12, 13], and further suggest that at least some degraders coexist with the secondary consumers. This result open an avenue to identify facilitation in natural samples.

Indeed, the relevance of ecological interactions in the assembly of natural POM communities is largely unknown [44]. In some studies, selective and stochastic processes explained a small amount of the variance [41], pointing towards alternative explanations such as the importance of competition [45]. In our experiments, we found an increase of rare taxa in the facilitation phase, which is likely explained by an increase in the number of microniches promoted by the degradation of the substrate, i.e. by facilitation interactions. Since facilitation might make these communities more persistent against perturbations, hence promoting diversity [46], our interpretation is consistent with the more robust biogeographic patterns observed for rare taxa [41, 47].

Next, we asked if it was possible to find distinctive traits for community-classes identified at the selection and facilitation phases. We analysed metagenomics sequencing data and metagenomics predictions generated from 16S sequences with PICRUSt [17]. In both cases, the emerging qualitative picture pointed towards a clear distinction between communities with higher motility, uptake of nutrients, chemotaxis and possibly high growth rates at the selection phase, and communities with a wide array of metabolic capabilities at late phases. Importantly, these patterns emerged after aggregating experiments that considered different substrates, and hosting different community compositions. Our results are hence aligned with previous observations pointing toward a decoupling between taxonomy and function [48], here represented by the distinct ecological traits of early and late colonizers.

We also considered the propensity of isolated strains to be found at different phases and analysed their genomes. In this analysis, we were able to identify a group of Vibrionaceae species with a high propensity to colonize in the attachment phase (<12h). This group showed a remarkably high proportion of ABC transporters related to the uptake of sugars, consistent with metagenomic predictions. For the comparison of isolates classified in the selection and facilitation phases we found a picture similar to the one found in the metagenomics analysis with, however, a lower significance.

The overarching picture emerging from our analysis is one in which mainly two types of strategies recapitulate the ecological succession, resembling the classical distinction between r- and K-selection. MacArthur envisioned this distinction in the colonization of islands in which, at the beginning of the colonization, there are unexploited abundant resources while, at later phases, resources are scarce and competition becomes harsh [18]. Species then experience different selective pressures at different phases, promoting the emergence of specialized strategies on the different scenarios, with r-strategists dominating earlier phases of colonization and K-strategists late phases. Although an oversimplification, the similarity of this picture with the one we observe in the colonization of synthetic particles in our experiments is remarkable. This picture has been criticized arguing that many examples presented as support of r/K-strategies can be explained by other mechanisms, in particular life-history traits [20]. This criticism is pertinent in macroscopic ecology, and it may be relevant for some species in our experiments (e.g. for those generalists appearing at both selection and facilitation phases), but we observe a global compositional turnover justifying this distinction.

In microbial ecology the r/K distinction is still used as a conceptual framework or as a postulate, due to a lack of mechanistic explanations connecting traits and ecological processes [21]. We provide some evidence to ground such mechanistic explanation. At very early time points (attachment phase), organisms encoding transporters related to the uptake of sugars colonize the particles. Also in this and in the next phase (selection) we observe a large proportion of genes related to flagellar assembly and chemotaxis. Apart from motility, the bacterial flagella are important for transient surface attachment [49]. Chemotaxis promotes migration along gradients of chemoattractants, which are created by the phycosphere of marine algae, decaying organic matter, and other biotic and abiotic sources [50]. It also enhances the expansion of motile bacteria towards uncolonized nutrient environments [51], a trait that is important for foraging among marine particles [43]. We also observed a high proportion of genes related to ribosome synthesis in the selection phase, which are characteristic of fast-growing bacteria [25, 26]. Therefore, bacteria found at the attachment and selection phases have traits to detect and to move towards abundant source of nutrients, with fast uptake and growth, a feast-and-famine strategy characteristic of copiotroph species [9].

Late colonizers exhibit a wide array of traits related with metabolism, suggesting that they experience a more competitive environment with more cells including more rare species and less abundant resources, characteristic of K-strategists. Analysing some pathways in detail, we found evidence for the existence of a trophic chain centered on the exchange of nitrogen-containing biomolecules including BCAA and ammonia, with early colonizers producing reduced forms of nitrogen and BCAA that are then taken and further degraded by late colonizers. Microbial degraders excrete amino acids during growth on digested polysaccharide [13]. Amino acids are thus likely to be part of the metabolites available to cells in the microenvironment of polysaccharide particles colonized by degraders. Interestingly, although late colonizers are also copiotrophs, some of the pathways found at late phases are characteristic of model representatives of oligotrophic bacteria such as Sphingopyxis alaskensis [9]. More specifically, in the metagenomes we found a high proportion of genes related with fatty acid biosynthesis or the degradation of xenobiotics like benzoate and aminobenzoate, consistent with a scenario in which carbon sources are more scarce.

In summary, our work provides a comprehensive analysis emphasizing the value of “domesticated” communities, namely growing natural communities under synthetic conditions, to gather insights into the selective forces structuring natural communities in complex processes such as ecological succession. Since, in natural environments, resources are often incorporated in bacterial niches not continuously but in periodic or random pulses (e.g. rain and drought in soil communities, food intake in gut microbes, marine snow in the ocean), we believe that the picture presented here in which a turnover in life-strategies recapitulate the ecological succession might be a general one, as also suggested in other studies [36, 52, 53]. This may thus be an important simplification to better understand and eventually predict microbial dynamics in the wild.

## Materials and Methods

### 16S amplicon sequencing

16S amplicon sequencing data was collected as described in [12, 13]. Amplicon sequencing data for samples collected from chitin, carageenan, agarose, alginate, and agarose-alginate hybrid particles are previously reported [12, 13]. Amplicon sequencing data for samples of seawater-associated microbes, chitosan particles, agarose-carrageenan, and agarose-chitosan particles are new to this study.

For amplicon sequencing samples reported in this study, 800 mL samples of coastal surface water collected in 2015 from Nahant, MA, USA were incubated with 100 particles/mL of a single particle type in triplicate in 1 L flasks. Particles were fabricated as described previously [13]. Bottles were sealed, and rotated end-over-end at room temperature. At 0, 12, 24, 36, 48, 60, 72, 108, 132, 156 and 204 h intervals, flasks were opened and 10 mL of bead/seawater mix was removed. Beads contained a magnetic core, and were separated from seawater using a neodymium magnet. Particle samples were resuspended twice in artificial seawater, and then stored for DNA extraction and sequencing library preparation. As described in detail previously [13], DNA was extracted from samples using a MasturePure extraction kit (Lucigen), and 16s rRNA amplicon libraries were prepared with primers 515F and 806R, which amplify the V4 region of the 16S rRNA gene.

Denoising was performed creating a parametric error model from a random set of 2M sequences, and this model was then used to identify erroneous sequence variants that were combined with the sequence variant that most likely originated following the pipeline implemented in the R Bioconductor dada2 package [54]. Functions from this package were also used for merging paired-end reads, trimming primer sequences, and dereplicating reads.

BarPlots and diversity analysis were performed with Phyloseq [55] after rarefying samples to the size of the sample with minimum number of reads (1K). Additional interactive visualizations of the ESVs trajectories clustered into Operational Taxonomic Units at different taxonomic thresholds were made with qiime2 [56], and are provided as Suppl. Materials. Ordination analysis were conducted computing the Bray-Curtis dissimilarity and then the dimensionality reduced with principal coordinate analysis (PCoA) with the R package Vegan [57].

### Metagenomes

Metagenomic sequencing libraries were prepared using genomic DNA extracted from particle samples at 0, 12, 24, 36, 48, 60, 72, 108, 132, 156 and 204 h timepoints. 2×250 paired-end sequencing libraries were prepared with Illumina Nexterra-XT adapters, using a modifications to the protocol previously validated for low DNA input [58].

Sequences were processed following the pipeline implemented in the Metagenomics Rapid Annotation using Subsystems Technology (MG-RAST) server version 4.0.3 [59]. In brief, the pipeline remove adapters with Skewer [60] and performs adapter clipping with fastq-mcf [61]. It then removes duplicated reads and assess the sequencing quality with DRISEE [62] and identifies potential contamination with Bowtie2 [63]. Functional annotations were obtained performing a de novo coding-region prediction with FragGeneScan [64], and translating the hits into amino-acid sequences. Protein sequences were clustered with cd-hit [65] and a sequence similarity search was performed with BLAT [66] against the M5NR database [67], considering a an e-value cut-off of 10^−5^. minimum identity cut-off of 60% and minimum length of sequence alignment of 15 nucleotides. KEGG annotations were considered for donwstream analysis. Samples with less than 10^3^ annotated genes (approximately corresponding to those with less than 10^4^ reads) were discarded to prevent biases. This excluded from the analysis most samples belonging to the attachment phase.

### Strains isolation and sequencing

Isolation of taxa from marine particles, and sequencing of isolate genomes to create draft assemblies was performed as described previously [13]. Briefly, 36, 84, 156h particles were sampled, washed, and diluted 1:1. 1:10, and 1:100 into artificial seawater (Sigma S9883). Particles in each dilution were vortexed for 20 s to dislodge cells, and plated onto 1.5 Bacto agar plates made with Marine Broth (MB, Difco 2216) as a nutrient source, or with Tibbles-Rawling defined minimal media [12] with 0.05 % w/v low viscosity alginate (Sigma A1112), 0.04 % carrageenan (Sigma C1013), 0.1 % glucosamine, or of with only agar. Plates were incubated for 2 days at room temperature, after which colonies were picked from each plate and struck for isolation on fresh MB agar. Isolate purity and taxonomic identification was assessed by Sanger dideoxy sequencing of the full 16S rRNA gene using primers 8F and 1492R [68]. Pure cultures were stored in 25 % glycerol at −80, prior to revival for DNA extraction for sequencing library preparation. Draft genomes were sequenced by preparing 2×250 paired-end Illumina sequencing libraries using Nexterra-XT library preparation and indexing kits. Sequencing was performed on an Illumina HiSeq 2500 at the Whitehead Institute for Biomedical Research in Cambridge, MA, USA. Reads were trimmed and filtered to remove unpaired and low quality reads, and then assembled de novo into contigs using CLC Genomics Workbench version 11. Completeness and other assembly statistics were assessed using CheckM [69], and taxonomic identification was made using gtDBK [70]. Contigs were annotated using the RAST pipeline [71].

### Determination of community classes

We performed an unsupervised classification of the samples by fitting the communities abundances to a linear combination of Dirichlet distributions [15], avoiding biases attributed to distance-based clustering methods [72]. Considering a combination of distributions provides a flexible framework to fit subsets of communities to distributions with different parameters, hence improving the fit. To prevent overfitting, a penalization was considered for having more distributions in the combination by taking the one that maximizes the posterior evidence of the fit [15]. Subsets of communities contributing to the same Dirichlet distribution were considered to belong to the same community-class, which, in the context of neutral theory is interpreted as being drawn from the same metacommunity [14]. We additionally verified that all communities contributed to a single Dirichlet component (i.e. that they are unambiguously classified in the same community-class).

### Test of neutrality

We tested if the communities were compatible with a neutral assembly following the method presented in Ref. [14]. This method represents a major advance to test for neutrality in large communities, which would not be possible to address following the model proposed by Hubbel [73]. Briefly, the method follows a Bayesian approach which independently fits the matrix **X** describing the abundances of ESVs in samples belonging to a given community-class (our proxy of metacommunity), to a Hierarchical Dirichlet Process with speciation parameter *θ*, metacommunity’s distribution 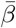 and immigration rates *I_i_* (*i* = 1,…, *M*), with *M* the number of samples in the class. Posterior samples of parameters, labelled with the index *k*, i.e. 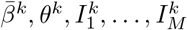, are then obtained. Next, synthetic matrices 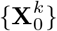 with the same number of samples and abundances than the class under study were sampled from each set of parameters *k*, simulating a Hierarchical Dirichlet Process (HDP). Finally, the metacommunity distribution parameters of each synthetic matrix are inferred 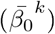. The method allows for the generation of both neutral metacommunities and neutral local communities, or of only neutral local communities (local model, hence with metacommunities being non-neutral). To test for neutrality, 50000 realizations of posterior samples were generated with 25000 being discarded as burn-in, and then 2500 selected among the last 25000 in intervals of 10 steps. The test considers that thee observed data appears neutral if the proportion of the log-likelihoods 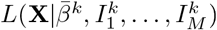 estimated from the observed data exceeds the correspondent log-likelihood estimated with synthetic data 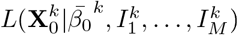 by some amount. This proportion is an empirical (pseudo) p-value and we considered neutrality rejected if *p* < 0.001. Computations were performed using the code provided in Ref. [14].

### Phylogenetic turnover and selection

To investigate the phylogenetic turnover through the ecological succession for each experiment, we computed a Mantel correlogram showing the relationship between community structure and time over different temporal distance classes. We considered significant tests with an adjusted p-value < 0.05. For each experiment, we rarefied the samples to 1000 reads and created a multiple sequence alignment with DECIPHER [74], and a phylogenetic tree with Neighbour Joining (R package ape [75]). The Mantel tests evaluated the correlation between the Unifrac distance [22] of the communities and their temporal distance (R package Vegan [57]).

We investigated different hypothesis regarding the influence of the environment in selecting taxonomically similar communities or in their taxonomic divergence following the analysis proposed in Ref. [24, 16]. We computed, for each pair of communities *k* and *l*, the weighted *β*-Mean Nearest Taxon Distance, defined as

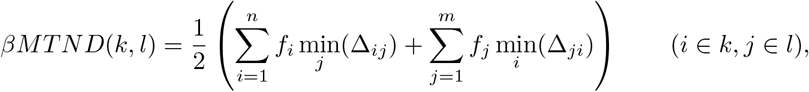

where *i* and *j* label ESVs, *f_i_* is the relative abundance of ESV *i*, and min_*j*_(Δ_*ij*_) is the minimum phylogenetic distance between the ESV *i* belonging to community *k* and all ESVs *j* in community *l*. In Ref. [24] it was shown that a *β*MNTD value significantly lower than the one obtained with the null model is expected when the environment constraints the communities composition. On the other hand, a value significantly higher than the null expectation would be found when different environmental conditions lead to divergent compositions among the communities. Other scenarios such as drift, would be expected for non-significant values. As a null model, we computed the *βMNTD* shuffling the ESVs in the nodes of the phylogenetic tree. We computed the null mean 〈*βMNTD*_rnd_〉 and standard deviation *σ*(*βMNTD*_rnd_) with 999 realizations of the null model, and then estimated the significance with a z-score (termed *β*–Nearest Taxon Index)[16]:

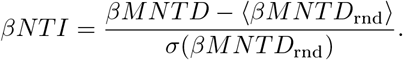

Those pairs of communities values fulfilling abs(**β*NTI*) > 2 were considered significant. Computations were conducted with R packages picante [76] and iCAMP [77].

### Predicted functional profiles

We predicted functional profiles from the ESVs found in 16S rRNA amplicon sequencing experiments. First, to reduce the number of ESVs considered we filtered those ESVs not showing a significant deviation in their abundances among the different substrates, or among the different time-points within any of the substrates, since it is unlikely they will contribute to differences in the functional profiles. We then developed a pipeline to investigate different hypothesis using DESeq2 [78] and Individual Value Analysis (R package indicspecies [79]).

We test preferences for specific substrates with a single model containing all beads’ samples and then looked for significant differences in the ESVs abundances between each pair of substrates (adjusted p-value < 0.01). We proceeded similarly to test preferences for particular times, now creating one model for each substrate individually, and then looking at significant differences between time points. A similar approximation was followed using Individual Value Analysis. Following these procedures we identified 2880 ESVs whose abundances are significantly higher in at least one substrate or time-point with any of the two methods. Considering these ESVs, the sparseness of the data set was reduced one order of magnitude while still representing 87% of the total number of reads. The final matrices for the different substrates are provided as Suppl. Material.

The functional profiles of the ESVs selected were estimated with PICRUSt v1.1.4 [17]. Quantitative and qualitative validation of the predictions were conducted computing the NSTI score [17], and with correlations between the profiles found in the predictions and those derived from the shotgun metagenome experiments. We also considered an additional quantitative validation of specific pathways showing discrepancies (see Suppl. Results).

### Estimation of ecological preferences for the isolates

We considered 65 strains isolated from previous experiments [12, 13] to investigate their genomes. To investigate which ESVs were in correspondence with the 16S sequences of the isolated strains, we made a BLAST database for the ESVs with the *makeblastdb* tool provided by NCBI-BLAST db [80] with options –parse_seqids –dbtype nucl. Then, we matched ESVs sharing 100% sequence identity with the 16S sequences of the strains using *blastn* with options –perc_identity100 –qcov_hsp_perc 100 –outfmt 6. One isolated strain had no 100% match with any ESV and it was discarded for downstream analysis. We finally associated the ecological preferences found for the ESVs to the isolates. To estimate the ecological preferences of the ESVs we estimated, for each ESV, a zero-inflated negative binomial generalized linear model [81, 82]. We considered as a response variable the abundances of the ESVs across all substrates, replicates and time points in beads’ samples, and the phase in which the abundance was measured as a dependent variable, then adding as an offset the logarithm of the total abundances of the sample [81]. The models were fitted with the zeroinfl function in package pscl [83]. We considered as a reference factor level the attachment phase, and established a code describing the significance of each phase relative to the reference (see Suppl. Results). We finally clustered the ESVs using this code to determine the ecological strategies (Suppl. Fig. 12).

### Phylogenetic analysis of the isolates

A phylogeny for the isolates was built using CheckM [69]. Briefly, 43 marker genes within each genome were used to place isolates into a reference phylogeny. Placement within the reference tree was used to derive topology of phylogenetic placements, and to assign taxonomy.

### Data availability

The short read 16 s rRNA amplicon sequencing data (BioSample: SAMN11023523-SAMN11023755) were deposted in NCBI with the BioProject identifier PRJNA478695. The 65 genomes have been deposited in NCBI with the BioProject identifiers PRJNA414740 and PRJNA478695. NCBI BioSample identifiers, some processed data and Supplementary Results including ESVs bar-plot visualizations are available in Zenodo (DOI: 10.5281/zenodo.5608678).

## Acknowledgements

We thank Mary-Ann Moran, Zackary Landry, Ben Roller, Magdalena San Román and other members of the Simons Collaboration Principles of Microbial Ecosystems (PRiME) for useful discussions. We also thank Christopher Quince for technical advice with some of the analysis.

## References

[1] Hutchins DA, Fu F. Microorganisms and ocean global change. Nature microbiology. 2017;2(6):1–11.

[2] Karl DM. Nutrient dynamics in the deep blue sea. Trends in Microbiology. 2002;10(9):410–418. doi:10.1016/S0966-842X(02)02430-7.

[3] Azam F, Malfatti F. Microbial structuring of marine ecosystems. Nature Reviews Microbiology. 2007;5(10):782–791. doi:10.1038/nrmicro1747.

[4] Lee C, Wakeham S, Arnosti C. Particulate Organic Matter in the Sea: The Composition Conundrum. AMBIO: A Journal of the Human Environment. 2004;33(8):565–575. doi:10.1579/0044-7447-33.8.565.

[5] Simon HM, Smith MW, Herfort L. Metagenomic insights into particles and their associated microbiota in a coastal margin ecosystem. Frontiers in Microbiology. 2014;5. doi:10.3389/fmicb.2014.00466.

[6] Polz MF, Cordero OX. Bacterial evolution: Genomics of metabolic trade-offs. Nature Microbiology. 2016;1(11):16181. doi:10.1038/nmicrobiol.2016.181.

[7] Bidle KD, Fletcher M. Comparison of free-living and particle-associated bacterial communities in the Chesapeake bay by stable low-molecular-weight RNA analysis. Applied and Environmental Microbiology. 1995;61(3):944–952.

[8] D’Ambrosio L, Ziervogel K, MacGregor B, Teske A, Arnosti C. Composition and enzymatic function of particle-associated and free-living bacteria: a coastal/offshore comparison. The ISME Journal. 2014;8(11):2167–2179. doi:10.1038/ismej.2014.67.

[9] Lauro FM, McDougald D, Thomas T, Williams TJ, Egan S, Rice S, et al. The genomic basis of trophic strategy in marine bacteria. Proceedings of the National Academy of Sciences. 2009; p. pnas.0903507106. doi:10.1073/pnas.0903507106.

[10] Bergauer K, Fernandez-Guerra A, Garcia JAL, Sprenger RR, Stepanauskas R, Pachiadaki MG, et al. Organic matter processing by microbial communities throughout the Atlantic water column as revealed by metaproteomics. Proceedings of the National Academy of Sciences. 2018;115(3):E400–E408. doi:10.1073/pnas.1708779115.

[11] Boeuf D, Edwards BR, Eppley JM, Hu SK, Poff KE, Romano AE, et al. Biological composition and microbial dynamics of sinking particulate organic matter at abyssal depths in the oligotrophic open ocean. Proceedings of the National Academy of Sciences. 2019;116(24):11824–11832. doi:10.1073/pnas.1903080116.

[12] Datta MS, Sliwerska E, Gore J, Polz MF, Cordero OX. Microbial interactions lead to rapid micro-scale successions on model marine particles. Nature Communications. 2016;7:11965.

[13] Enke TN, Datta MS, Schwartzman J, Cermak N, Schmitz D, Barrere J, et al. Modular assembly of polysaccharide-degrading marine microbial communities. Current Biology. 2019;29(9):1528–1535.

[14] Harris K, Parsons TL, Ijaz UZ, Lahti L, Holmes I, Quince C. Linking statistical and ecological theory: Hubbell’s unified neutral theory of biodiversity as a hierarchical Dirichlet process. Proceedings of the IEEE. 2015;.

[15] Holmes I, Harris K, Quince C. Dirichlet multinomial mixtures: generative models for microbial metagenomics. PloS One. 2012;7(2):e30126.

[16] Stegen JC, Lin X, Fredrickson JK, Chen X, Kennedy DW, Murray CJ, et al. Quantifying community assembly processes and identifying features that impose them. The ISME journal. 2013;7(11):2069–2079.

[17] Langille MG, Zaneveld J, Caporaso JG, McDonald D, Knights D, Reyes JA, et al. Predictive functional profiling of microbial communities using 16S rRNA marker gene sequences. Nature Biotechnology. 2013;31(9):814–821.

[18] Pianka ER. On r-and K-selection. The American Naturalist. 1970;104(940):592–597.

[19] MacArthur RH, Wilson EO. The theory of island biogeography. Princeton university press; 1967.

[20] Reznick D, Bryant MJ, Bashey F. r-and K-selection revisited: the role of population regulation in life-history evolution. Ecology. 2002;83(6):1509–1520.

[21] Andrews JH, Harris RF. r-and K-selection and microbial ecology. In: Advances in Microbial Ecology. Springer; 1986. p. 99–147.

[22] Lozupone C, Lladser ME, Knights D, Stombaugh J, Knight R. UniFrac: an effective distance metric for microbial community comparison. The ISME journal. 2011;5(2):169–172.

[23] Wang J, Shen J, Wu Y, Tu C, Soininen J, Stegen JC, et al. Phylogenetic beta diversity in bacterial assemblages across ecosystems: deterministic versus stochastic processes. The ISME journal. 2013;7(7):1310–1321.

[24] Dini-Andreote F, Stegen JC, van Elsas JD, Salles JF. Disentangling mechanisms that mediate the balance between stochastic and deterministic processes in microbial succession. Proceedings of the National Academy of Sciences. 2015;112(11):E1326–E1332.

[25] Klappenbach JA, Dunbar JM., Schmidt TM. rRNA Operon Copy Number Reflects Ecological Strategies of Bacteria. Applied and Environmental Microbiology. 2000;66(4):1328–1333. doi:10.1128/AEM.66.4.1328-1333.2000.

[26] Roller BR, Stoddard SF, Schmidt TM. Exploiting rRNA operon copy number to investigate bacterial reproductive strategies. Nature Microbiology. 2016;1:16160.

[27] Cui J, Davidson AL. ABC solute importers in bacteria. Essays in Biochemistry. 2011;50:85–99. doi:10.1042/bse0500085.

[28] Shimada T, Kori A, Ishihama A. Involvement of the ribose operon repressor RbsR in regulation of purine nucleotide synthesis in Escherichia coli. FEMS Microbiology Letters. 2013;344(2): 159–165. doi:10.1111/1574-6968.12172.

[29] D’Arrigo I, Cardoso JGR, Rennig M, Sonnenschein N, Herrgård MJ, Long KS. Analysis of Pseudomonas putida growth on non-trivial carbon sources using transcriptomics and genome-scale modelling. Environmental Microbiology Reports. 2019;11(2):87–97. doi:10.1111/1758-2229.12704.

[30] Nikata T, Sakai Y, Shibata K, Kato J, Kuroda A, Ohtake H. Molecular analysis of the phosphate-specific transport (pst) operon ofPseudomonas aeruginosa. Molecular and General Genetics MGG. 1996;250(6):692–698. doi:10.1007/BF02172980.

[31] Ogura M. ZnuABC and ZosA zinc transporters are differently involved in competence development in Bacillus subtilis. The Journal of Biochemistry. 2011;150(6):615–625. doi:10.1093/jb/mvr098.

[32] Vondenhoff GHM, Blanchaert B, Geboers S, Kazakov T, Datsenko KA, Wanner BL, et al. Characterization of Peptide Chain Length and Constituency Requirements for YejABEF-Mediated Uptake of Microcin C Analogues. Journal of Bacteriology. 2011;193(14):3618–3623. doi:10.1128/JB.00172-11.

[33] Massey LK, Sokatch JR, Conrad RS. Branched-chain amino acid catabolism in bacteria. Bacteriological Reviews. 1976;40(1):42–54.

[34] Pascual-García A. Phylogenetic Core Groups: a promising concept in search of a consistent methodological framework. Microbiome. 2021;9(1):1–9.

[35] Arumugam M, Raes J, Pelletier E, Le Paslier D, Yamada T, Mende DR, et al. Enterotypes of the human gut microbiome. Nature. 2011;473(7346):174–180.

[36] Pascual-García A, Bell T. Community-level signatures of ecological succession in natural bacterial communities. Nature communications. 2020;11(1):1–11.

[37] Ramoneda J, Hawes I, Pascual-García A, J Mackey T, Y Sumner D, D Jungblut A. Importance of environmental factors over habitat connectivity in shaping bacterial communities in microbial mats and bacterioplankton in an Antarctic freshwater system. FEMS Microbiology Ecology. 2021;97(4):fiab044.

[38] Caruso T, Chan Y, Lacap DC, Lau MCY, McKay CP, Pointing SB. Stochastic and deterministic processes interact in the assembly of desert microbial communities on a global scale. The ISME Journal. 2011;5(9):1406–1413. doi:10.1038/ismej.2011.21.

[39] Dumbrell AJ, Nelson M, Helgason T, Dytham C, Fitter AH. Relative roles of niche and neutral processes in structuring a soil microbial community. The ISME Journal. 2009;4(3):337–345. doi:10.1038/ismej.2009.122.

[40] Sunagawa S, Coelho LP, Chaffron S, Kultima JR, Labadie K, Salazar G, et al. Structure and function of the global ocean microbiome. Science. 2015;348(6237):1261359.

[41] Wu W, Logares R, Huang B, Hsieh Ch. Abundant and rare picoeukaryotic sub-communities present contrasting patterns in the epipelagic waters of marginal seas in the northwestern Pacific Ocean. Environmental Microbiology. 2017;19(1):287–300. doi:https://doi.org/10.1111/1462-2920.13606.

[42] Vergin KL, Jhirad N, Dodge J, Carlson CA, Giovannoni SJ. Marine bacterioplankton consortia follow deterministic, non-neutral community assembly rules. Aquatic Microbial Ecology. 2017;79(2):165–175.

[43] Fernandez VI, Yawata Y, Stocker R. A Foraging Mandala for Aquatic Microorganisms. The ISME Journal. 2019;13(3):563–575. doi:10.1038/s41396-018-0309-4.

[44] Liu J, Meng Z, Liu X, Zhang XH. Microbial assembly, interaction, functioning, activity and diversification: a review derived from community compositional data. Marine Life Science & Technology. 2019;1(1):112–128.

[45] Zhang Y, Zhao Z, Dai M, Jiao N, Herndl GJ. Drivers shaping the diversity and biogeography of total and active bacterial communities in the South China Sea. Molecular Ecology. 2014;23(9):2260–2274.

[46] Pascual-García A, Bonhoefier S, Bell T. Metabolically cohesive microbial consortia and ecosystem functioning. Philosophical Transactions of the Royal Society B. 2020;375(1798):20190245.

[47] Mo Y, Zhang W, Yang J, Lin Y, Yu Z, Lin S. Biogeographic patterns of abundant and rare bacterioplankton in three subtropical bays resulting from selective and neutral processes. The ISME journal. 2018;12(9):2198–2210.

[48] Louca S, Parfrey LW, Doebeli M. Decoupling function and taxonomy in the global ocean microbiome. Science. 2016;353(6305): 1272–1277.

[49] Kimkes TEP, Heinemann M. How bacteria recognise and respond to surface contact. FEMS Microbiology Reviews. 2020;44(1):106–122. doi:10.1093/femsre/fuz029.

[50] Stocker R, Seymour JR. Ecology and Physics of Bacterial Chemotaxis in the Ocean. Microbiology and Molecular Biology Reviews. 2012;76(4):792–812. doi:10.1128/MMBR.00029-12.

[51] Cremer J, Honda T, Tang Y, Wong-Ng J, Vergassola M, Hwa T. Chemotaxis as a navigation strategy to boost range expansion. Nature. 2019;575(7784):658–663. doi:10.1038/s41586-019-1733-y.

[52] Tipton L, Darcy JL, Hynson NA. A developing symbiosis: enabling cross-talk between ecologists and microbiome scientists. Frontiers in microbiology. 2019;10:292.

[53] Freilich S, Kreimer A, Meilijson I, Gophna U, Sharan R, Ruppin E. The large-scale organization of the bacterial network of ecological co-occurrence interactions. Nucleic Acids Research. 2010;38(12):3857–3868. doi:10.1093/nar/gkq118.

[54] Callahan BJ, McMurdie PJ, Rosen MJ, Han AW, Johnson AJA, Holmes SP. DADA2: High-resolution sample inference from Illumina amplicon data. Nature Methods. 2016;13(7):581–583. doi:10.1038/nmeth.3869.

[55] McMurdie PJ, Holmes S. phyloseq: an R package for reproducible interactive analysis and graphics of microbiome census data. PloS One. 2013;8(4):e61217.

[56] Caporaso JG, Kuczynski J, Stombaugh J, Bittinger K, Bushman FD, Costello EK, et al. QIIME allows analysis of high-throughput community sequencing data. Nature Methods. 2010;7(5):335–336.

[57] Oksanen J, Blanchet FG, Kindt R, Legendre P, Minchin P, O’hara R, et al. Community ecology package. R package version. 2013;2(0).

[58] Rinke C, Low S, Woodcroft BJ, Raina JB, Skarshewski A, Le XH, et al. Validation of picogram-and femtogram-input DNA libraries for microscale metagenomics. PeerJ. 2016;4:e2486.

[59] Keegan KP, Glass EM, Meyer F. MG-RAST, a metagenomics service for analysis of microbial community structure and function. In: Microbial environmental genomics (MEG). Springer; 2016. p. 207–233.

[60] Jiang H, Lei R, Ding SW, Zhu S. Skewer: a fast and accurate adapter trimmer for next-generation sequencing paired-end reads. BMC bioinformatics. 2014;15(1):1–12.

[61] Aronesty E. Comparison of sequencing utility programs. The open bioinformatics journal. 2013;7(1).

[62] Keegan KP, Trimble WL, Wilkening J, Wilke A, Harrison T, D’Souza M, et al. A platform-independent method for detecting errors in metagenomic sequencing data: DRISEE. PLoS Comput Biol. 2012;8(6):e1002541.

[63] Langmead B, Salzberg SL. Fast gapped-read alignment with Bowtie 2. Nature methods. 2012;9(4):357.

[64] Rho M, Tang H, Ye Y. FragGeneScan: predicting genes in short and error-prone reads. Nucleic acids research. 2010;38(20):e191–e191.

[65] Fu L, Niu B, Zhu Z, Wu S, Li W. CD-HIT: accelerated for clustering the next-generation sequencing data. Bioinformatics. 2012;28(23):3150–3152.

[66] Kent WJ. BLAT–the BLAST-like alignment tool. Genome research. 2002;12(4):656–664.

[67] Wilke A, Harrison T, Wilkening J, Field D, Glass EM, Kyrpides N, et al. The M5nr: a novel non-redundant database containing protein sequences and annotations from multiple sources and associated tools. BMC bioinformatics. 2012;13(1):1–5.

[68] Turner S, Pryer KM, Miao VP, Palmer JD. Investigating deep phylogenetic relationships among cyanobacteria and plastids by small subunit rRNA sequence analysis 1. Journal of Eukaryotic Microbiology. 1999;46(4):327–338.

[69] Parks DH, Imelfort M, Skennerton CT, Hugenholtz P, Tyson GW. CheckM: assessing the quality of microbial genomes recovered from isolates, single cells, and metagenomes. Genome research. 2015;25(7):1043–1055.

[70] Chaumeil PA, Mussig AJ, Hugenholtz P, Parks DH. GTDB-Tk: a toolkit to classify genomes with the Genome Taxonomy Database. Bioinformatics. 2019;36(6):1925–1927. doi:10.1093/bioinformatics/btz848.

[71] Aziz RK, Bartels D, Best AA, DeJongh M, Disz T, Edwards RA, et al. The RAST Server: rapid annotations using subsystems technology. BMC genomics. 2008;9(1):1–15.

[72] Warton DI, Wright ST, Wang Y. Distance-based multivariate analyses confound location and dispersion effects. Methods in Ecology and Evolution. 2012;3(1):89–101.

[73] Hubbell SP. The unified neutral theory of biodiversity and biogeography (MPB-32). Princeton University Press; 2001.

[74] Wright ES. Using DECIPHER v2.0 to analyze big biological sequence data in R. The R Journal. 2016;8(1):352–359.

[75] Paradis E, Schliep K. ape 5.0: an environment for modern phylogenetics and evolutionary analyses in R. Bioinformatics. 2019;35:526–528.

[76] Kembel SW, Cowan PD, Helmus MR, Cornwell WK, Morlon H, Ackerly DD, et al. Picante: R tools for integrating phylogenies and ecology. Bioinformatics. 2010;26:1463–1464.

[77] Ning D, Yuan M, Wu L, Zhang Y, Guo X, Zhou X, et al. A quantitative framework reveals ecological drivers of grassland microbial community assembly in response to warming. Nature Communications. 2020;doi:10.1038/s41467-020-18560-z.

[78] Love MI, Huber W, Anders S. Moderated estimation of fold change and dispersion for RNA-seq data with DESeq2. Genome Biology. 2014;15:550. doi:10.1186/s13059-014-0550-8.

[79] Cáceres MD, Legendre P. Associations between species and groups of sites: indices and statistical inference. Ecology. 2009;90(12):3566–3574.

[80] Altschul SF, Gish W, Miller W, Myers EW, Lipman DJ. Basic local alignment search tool. Journal of Molecular Biology. 1990;215(3):403–410.

[81] Xia Y, Sun J, Chen DG, et al. Statistical analysis of microbiome data with R. vol. 847. Springer; 2018.

[82] Xu L, Paterson AD, Turpin W, Xu W. Assessment and selection of competing models for zero-inflated microbiome data. PloS one. 2015;10(7):e0129606.

[83] Zeileis A, Kleiber C, Jackman S. Regression Models for Count Data in R. Journal of Statistical Software. 2008;27(8).

