## Supplementary Materials for "Turnover in life-strategies recapitulates marine microbial succession colonizing model particles"

### Supplementary Material
### Turnover in life-strategies recapitulates microbial succession in synthetic marine particles

Alberto Pascual-García[1,*], Julia Schwartzman[2], Arion Iffland-Stettner[1], Otto X. Cordero[2,♮] and Sebastian Bonhoeffer[1,♮].

November 5, 2021

(1) Institute of Integrative Biology, ETH-Zürich; Zürich, Switzerland
(2) Add MIT affiliation here
(♮) Equal contribution

### 1 Supplementary Figures

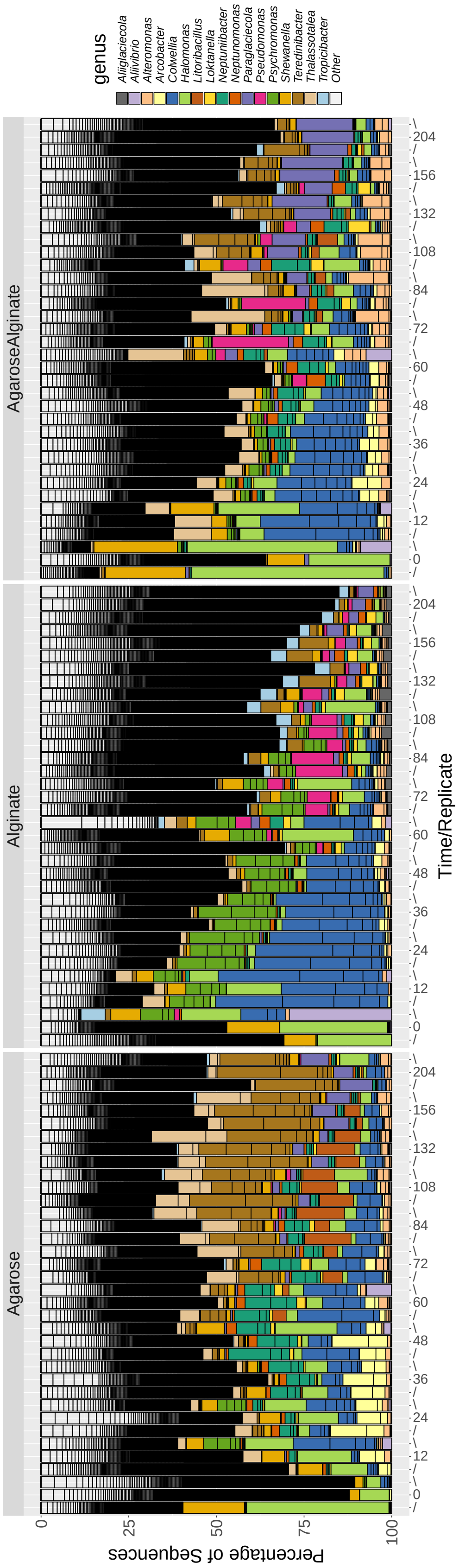

Figure 1: **Bar plots for attached populations in beads** of agarose, alginate and a mix of both agarose and alginate. Each time point embraces three replicates, showing a remarkable reproducibility. Genera among the 15 most abundant otus in at least one time point are highlighted with the remainder classified as Others. Figure reproduced from Ref. [1] with permission.

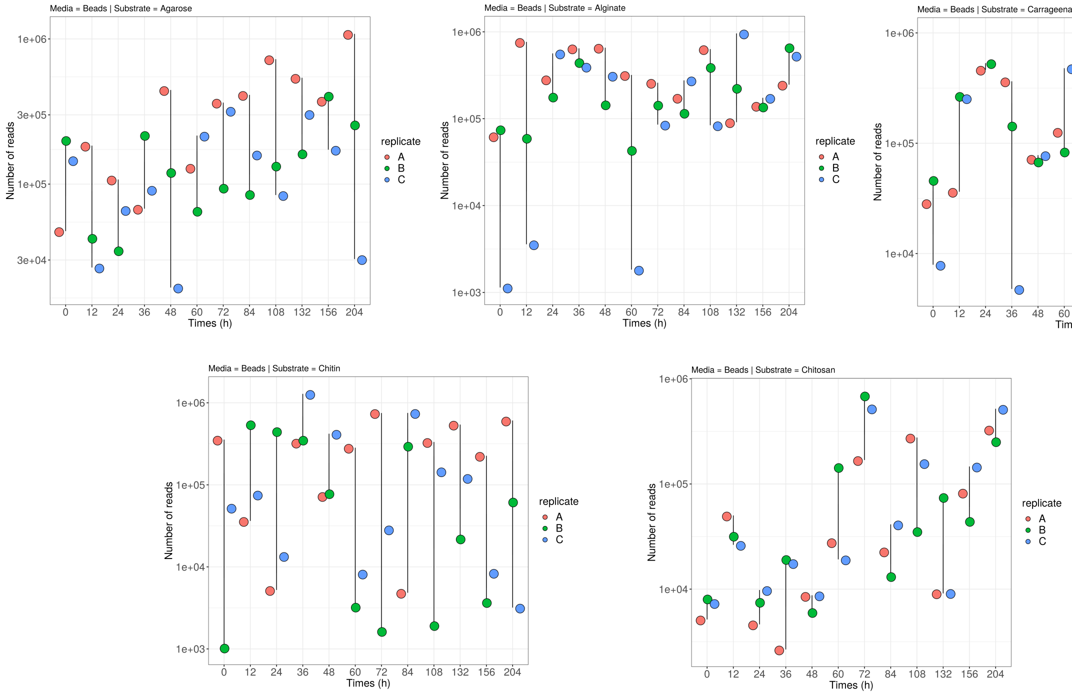

Figure 2: **Number of reads** for attached populations in the pure substrates. To help identifying each group of three replicates, a line was added on the replicate in the middle (B) connecting the maximum and minimum of the three replicates belonging to the same time-point.

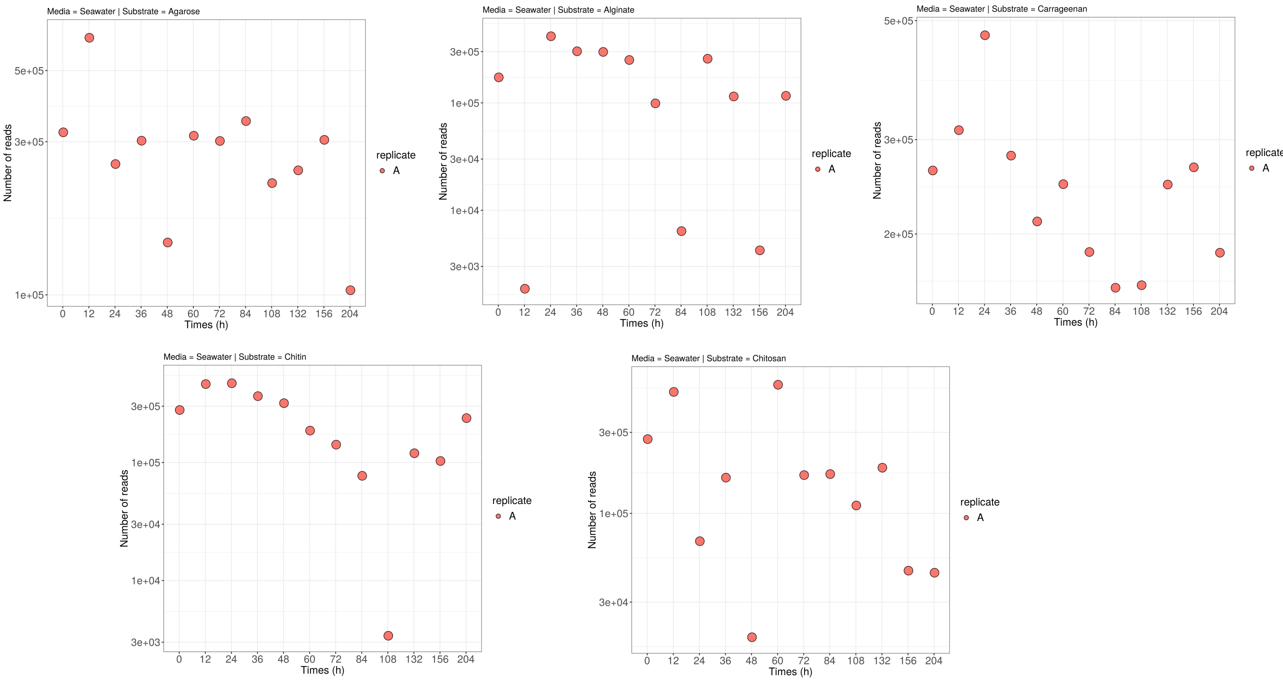

Figure 3: **Number of reads** for populations present in the beads' surrounding seawater.

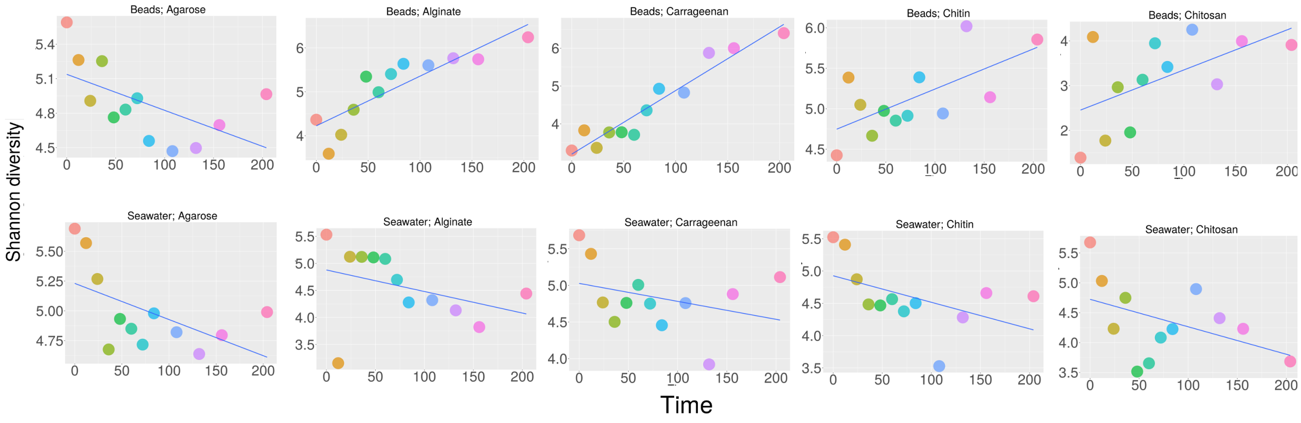

Figure 4: **Shannon diversity** for populations attached to beads (first row) and present in surrounding water (second row) for each substrate (columns) at the different time points.

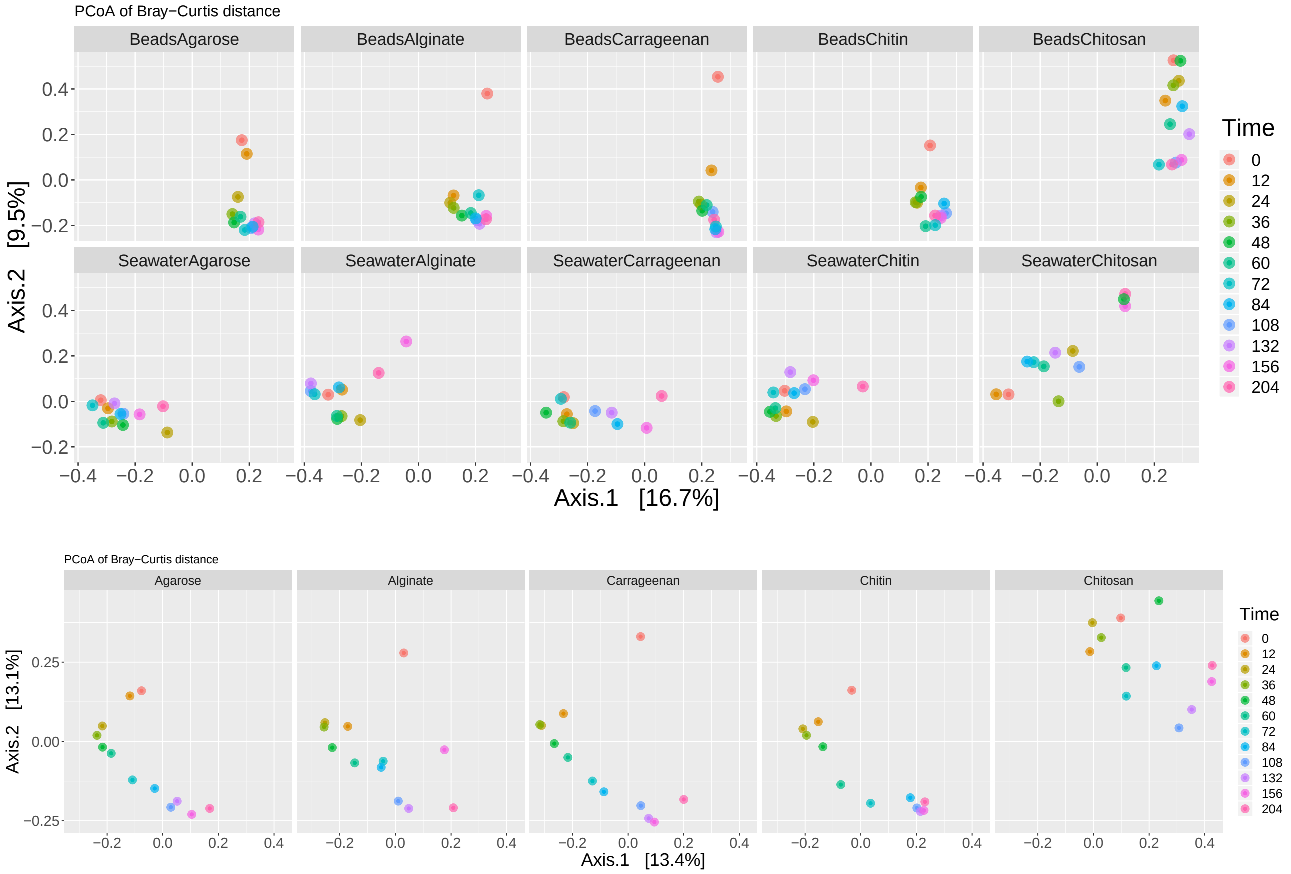

Figure 5: **Principal coordinate analysis (PCoA) (i)** of Bray-Curtis similarity between communities. (Top) Communities sampled from beads (first row) and seawater (second row) are clearly separated when projected in the reduced space, and they are more similar at latest times. For chitosan, the beads' trajectory reaches the bottom of axis 2, and the trajectory representing seawater samples evolves towards the positive quadrant of the figure. (Bottom) Considering only the trajectories of communities on the beads we observe their similarity irrespective of the substrate considered, suggesting that temporal dynamics have a more preeminent role than the specific substrate. Chitosan is again an exception, with a trajectory constrained to the positive quadrant, consistent with the idea that this substrate is more recalcitrant and remains in the region occupied by some of the other substrates only at the first time-point. For these representations the three replicates were aggregated.

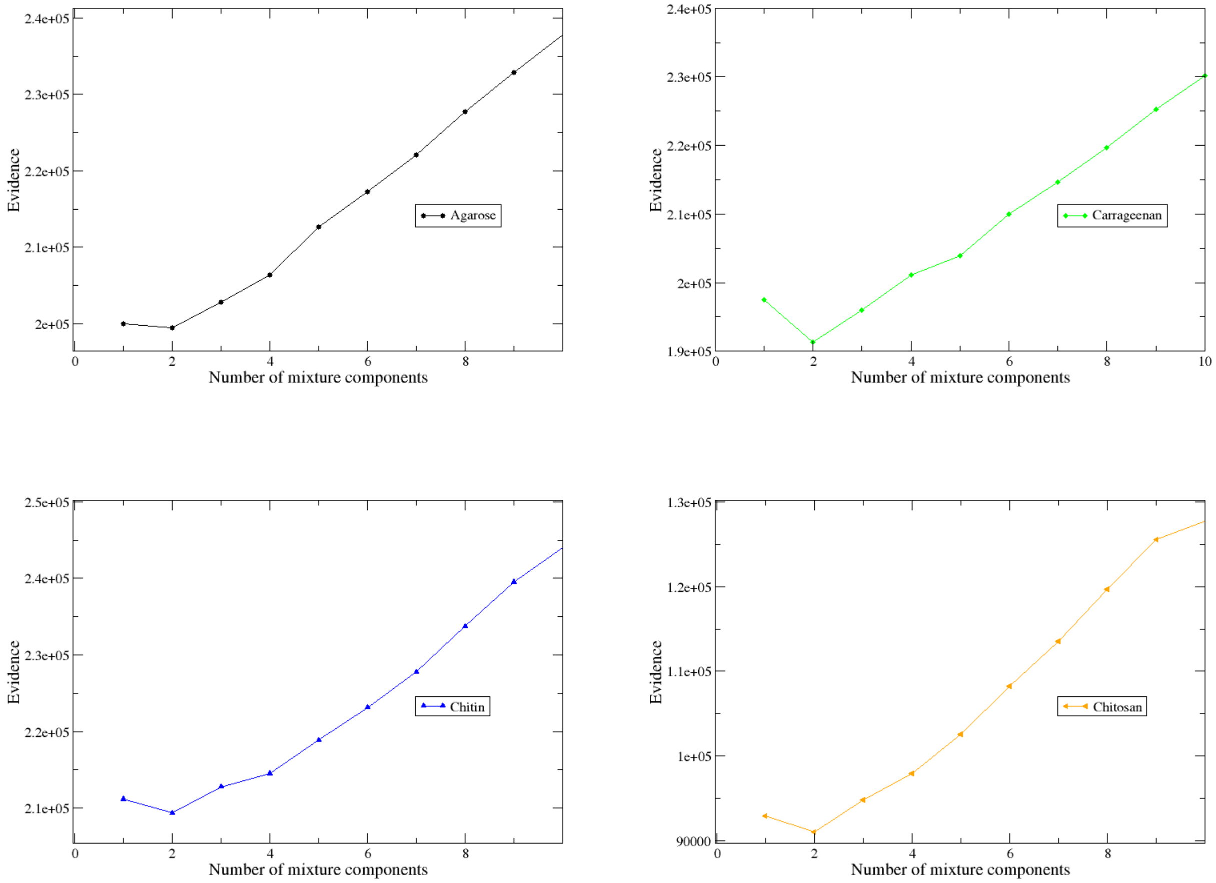

Figure 6: **Posterior evidence against the number of mixture components** used in the fit of data, for each substrate. The minimum indicates the optimal number of community classes.

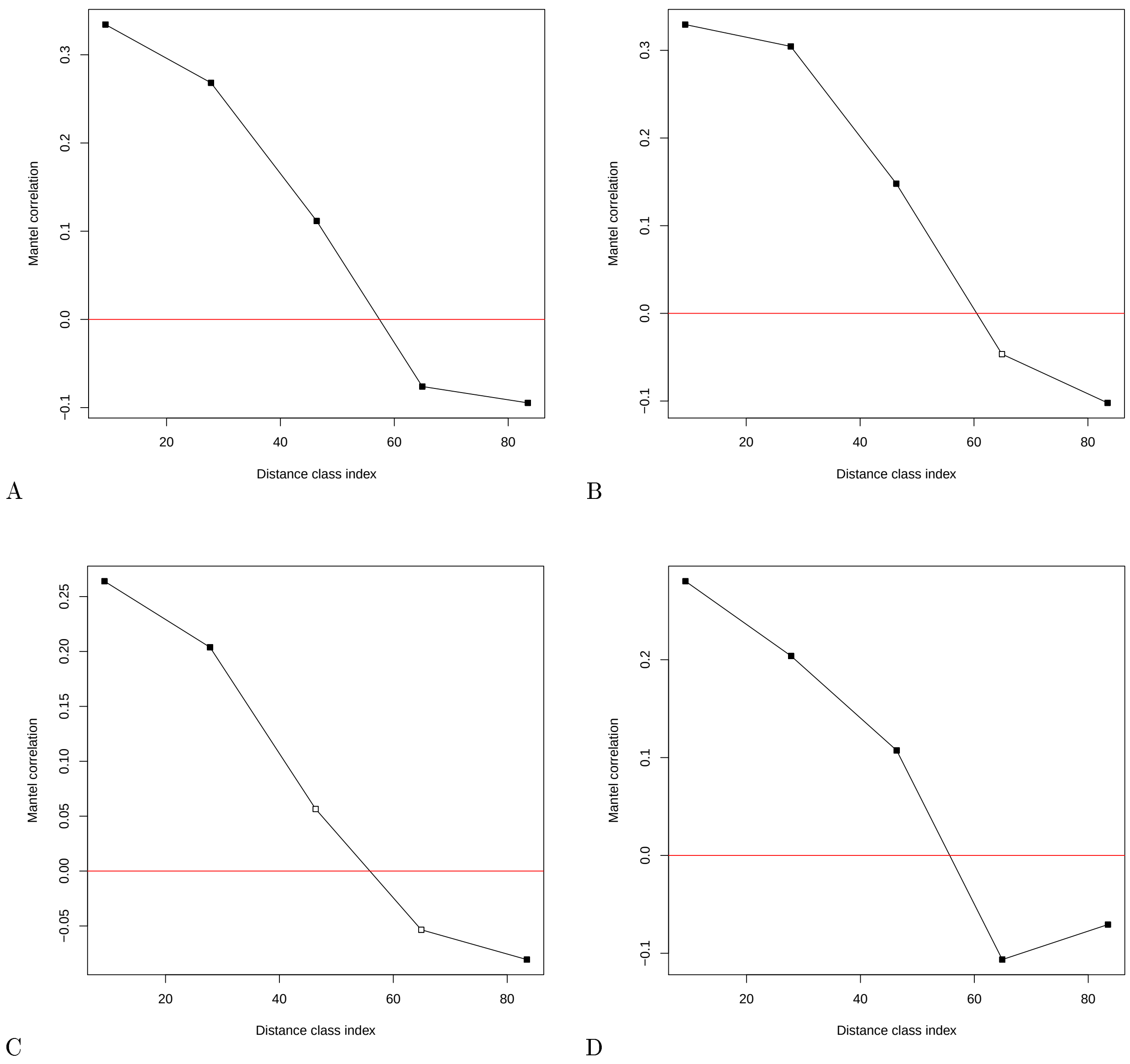

Figure 7: **Mantel correlogram against distance classes for pure substrates.** Correlation between the Unifrac distance and the temporal distance for samples classified at short distances (left-hand-side of the x-axis) or long distances (right-hand-side) for agarose (A), Carregeenan (B), Chitin (C) and Chitosan (D). Significant correlations are shown with filled boxes. Results for alginate are shown in the Main Text.

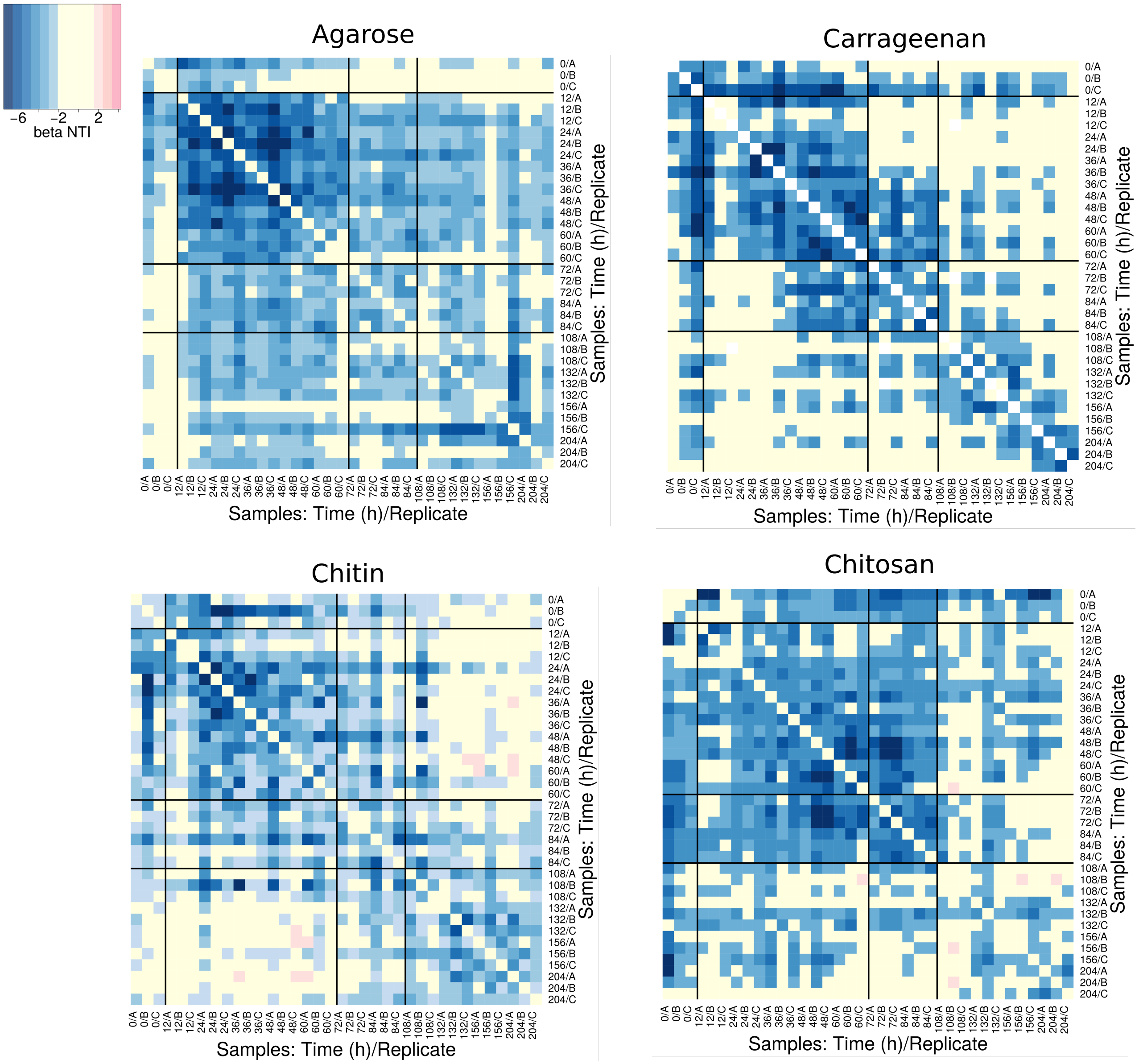

Figure 8: **All-against-all beta Nearest Taxon Index ($\beta NTI$)** similarity of communities within each experiment. Each heatmap represents one experiment and samples are ordered by time-point and replicate. Absolute values larger than two are considered significant.

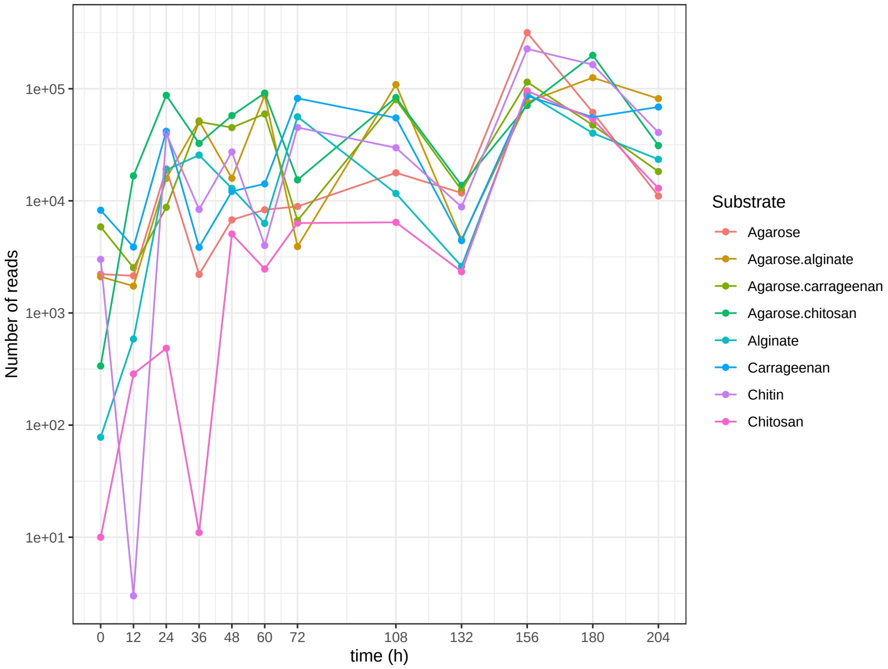

Figure 9: **Number of reads in metagenomes experiments.**

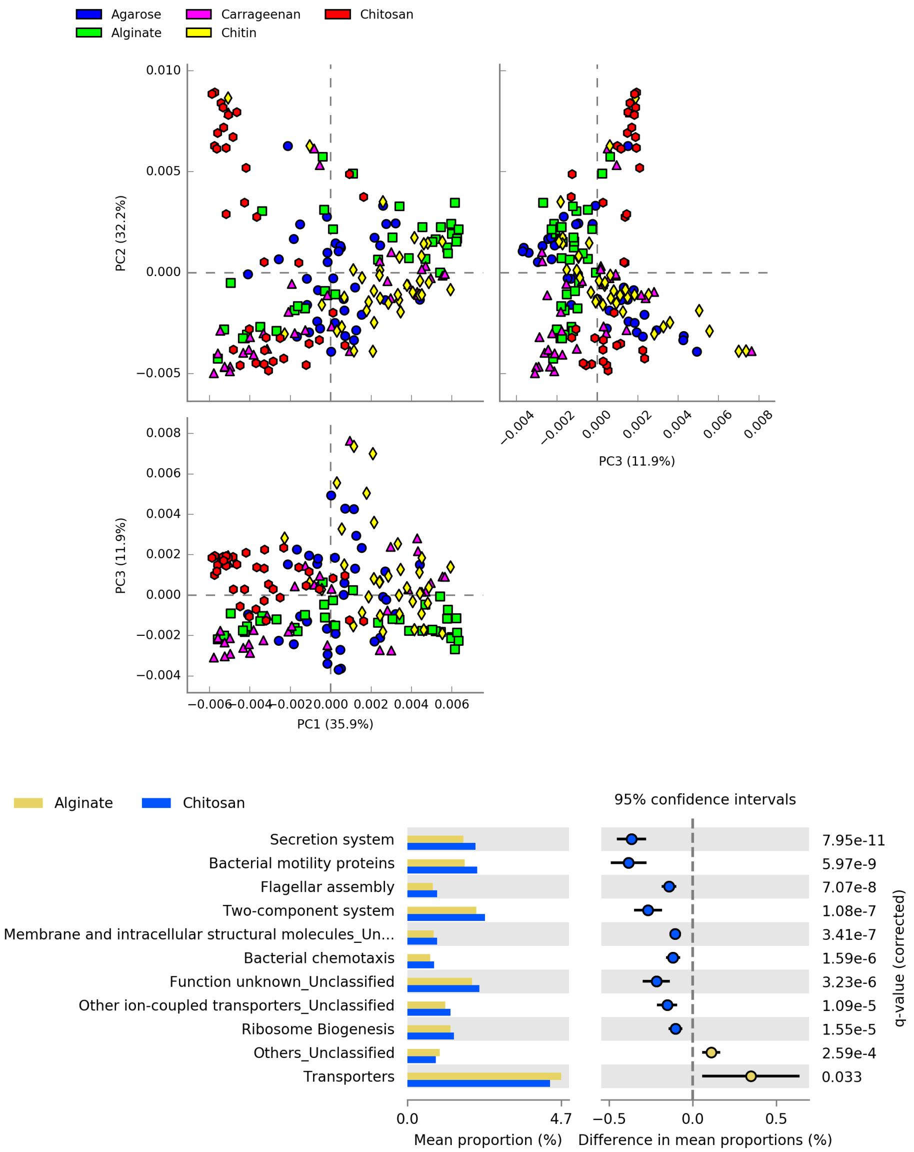

Figure 10: **Comparison of metagenomics predictions across substrates.**(Top) Principal component analysis of metagenomic predicted profiles in beads samples. Experiments with mixed substrates were removed for clarity. Contrary to the clear clustering corresponding to the different temporal stages shown in the PCA presented in the Main Text, by identifying the samples by their substrate does not lead to any clear clustering, with most of the substrates having dots spread in the three directions. An exception is chitosan, concentrating more points in the region where the attachment stage is, suggesting a slower dynamics. This is confirmed when we compare the mean proportion o genes in chitosan with respect to other substrates (e.g. with respect to alginate, Bottom figure), since we observe that the pathways more represented in chitosan are those we associated to the early-stages of the colonization when all substrates were aggregated together.

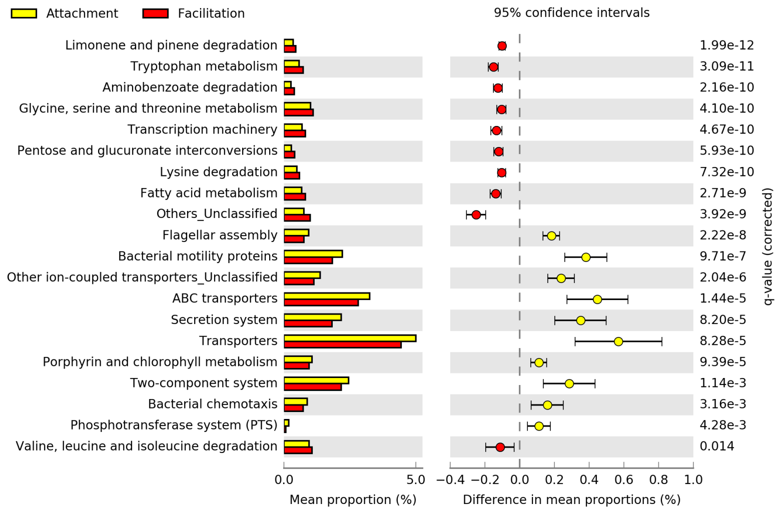

Figure 11: **Comparison of metagenomics predictions between attachment and facilitation stages.** Difference in the mean proportions of genes between communities at the selection and facilitation stages in metagenomic and in PiCRUST predictions. Each row in the diagram represents genes classified in the KEGG pathway indicated. The first column represents the mean proportions of the genes in the pathway for each stage, and the second column the difference between those proportions. Adjusted Bejamini-Hochberg p-values and 95% CI intervals are indicated. Pathways with effect sizes lower than 0.1 were filtered.

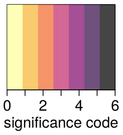

| Stage | Significance Code (SC) | Abundance |
|---|---|---|
| Selection (reference) | SC(Selection) = 0 | Not significantly > 0 |
| | SC(Selection) = 1 | Significantly > 0 (p-val < 0.05) |
| | SC(Selection) = 2 | Significantly > 0 (p-val < 0.01) |
| | SC(Selection)= 3 | Significantly > 0 (p-val < 0.001) |
| Others | SC = SC(Selection) | Similar to the selection stage |
| | SC = SC(Selection) ± 1 | Higher (+1), or smaller (-1) than in the selection stage (p-val < 0.05) |
| | SC = SC(Selection) ± 2 | Higher (+2), or smaller (-2) than in the selection stage (p-val < 0.01) |
| | SC = SC(Selection) ± 3 | Higher (+3), or smaller (-3) than in the selection stage (p-val < 0.001) |

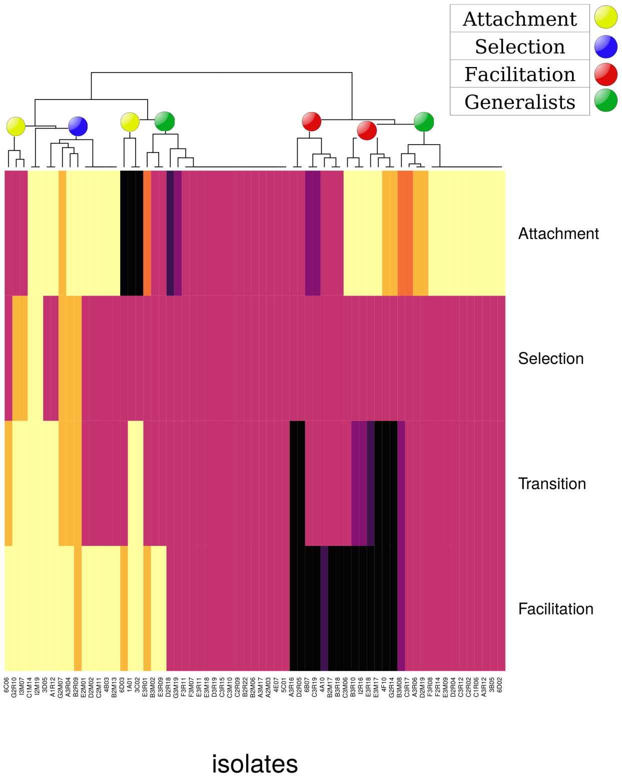

Figure 12: **Significance of the generalized linear model coefficients.** The heatmap shows the significance of the coefficients of a zero-inflated negative binomial model with the phases as predictors (rows) and the abundances of the ESVs matching 100% sequence identity the isolated strains (columns) as responses. In the Table it is indicated the meaning of the significance code: the selection phase is taken as a reference (intercept) and the value of the significance code ranges from zero (the abundance of the ESV is not significantly different than zero in that phase) to three (significantly positive, $p < 10^{-3}$). The significance code of the remainder phases is fixed relative to the selection phase. A higher (lower) value indicates that the abundance of the ESV in the phase indicated is significantly higher (lower) than the one observed at the selection phase, the more significant is the difference in abundances the larger is the difference in the code. The isolates are then clustered (top of the heatmap) and the clusters manually associated to specific stages (indicated with coloured circles in the dendrogram) after visual inspection.

| | Metagenomes | | Isolates | |
|---|---|---|---|---|
| | Experiments | PiCRUST | | |
| Pathway / Stages compared | Selection / facilitation | Selection / facilitation | Selection / facilitation | Attachment / facilitation |
| **Cellular processes** | | | | |
| *Cell motility* | | | | |
| Flagellar assembly | Selection | Selection | Selection (n.s.) | X |
| Bacterial motility proteins | X | Selection | X | X |
| Bacterial chemotaxis | Selection | Selection | Selection | X |
| *Cellular community* | | | | |
| Biofilm formation – E. coli | X | X | X | Attachment |
| **Human Diseases** | | | | |
| *Drug resistance* | | | | |
| beta-Lactam resistance | X | X | X | Attachment |
| **Genetic Information Processing** | | | | |
| *Transcription* | | | | |
| Transcription machinery | X | Facilitation | X | X |
| *Translation* | | | | |
| Aminoacyl-tRNA biosynthesis | Selection | X | X | Facilitation |
| Ribosome | Selection | X | X | X |
| Ribosome biogenesis | X | X | Selection (n.s.) | Attachment |
| *Replication and repair* | | | | |
| Homologous recombination | Selection | X | X | X |
| Mismatch repair | Selection | X | X | X |
| Chaperones and folding catalysts | X | Selection | Selection (n.s.) | Attachment |
| **Environmental Information Processing** | | | | |
| *Membrane transport* | | | | |
| ABC-transporters | Selection | X | X | Attachment |
| Phosphotransferase system (PTS) | X | X | X | Attachment |
| Transporters | X | X | X | Attachment |
| Bacterial secretion system | X | Selection | X | X |
| Secretion system | X | Selection | X | X |
| Signal transduction | | | | |
| Two-component system | X | Selection | Selection (n.s.) | X |
| *Signalling and cellular processing (unclassified)* | | | | |
| Structural proteins | X | X | X | Attachment |

Table 1: **Summary of KEGG pathways (i)** showing significant differences between phases in their mean proportions (rows) for the three datasets considered (columns). The stages compared in each dataset are indicated. Cells indicate the phase in which the mean proportion is significantly higher, having an "X" if no significant differences were found. Cells labelled as (n.s.) indicate that the difference is not significance when corrected for multiple testing. This table summarizes pathways classified in KEGG level 1 as Cellular Processes, Human Diseases, Genetic Information Processes and Environmental Information Processes, relatively higher for bacteria present in the selection and attachment phases.

| Pathway ▼ / Stages compared ► | Metagenomes: Experiments — Selection / facilitation | Metagenomes: PiCRUST — Selection / facilitation | Isolates — Selection / facilitation | Isolates — Attachment / facilitation |
|---|---|---|---|---|
| **Metabolism** | | | | |
| *Energy metabolism* | | | | |
| Nitrogen metabolism | X | Selection | X | X |
| Energy metabolism (unclassified) | X | X | X | Attachment |
| *Metabolism of cofactors and vitamins* | | | | |
| Thiamine metabolism | Selection | X | X | X |
| Porphyrin and chlorophyll metabolism | Selection | X | X | X |
| *Nucleotide metabolism* | | | | |
| Purine metabolism | X | X | X | Attachment |
| *Carbohydrate metabolism* | | | | |
| Citrate cycle (TCA) | X | Selection | X | X |
| Glyoxylate and dicarboxilate metabolism | X | X | Facilitation (n.s.) | Facilitation |
| Pentose phosphate pathway | X | Facilitation | X | X |
| Fructose and mannose metabolism | X | Facilitation | Facilitation (n.s.) | X |
| Pentose and glucuronate interconversions | Facilitation | Facilitation | Facilitation (n.s.) | X |
| Starch and sucrose metabolism | Facilitation | Facilitation | X | X |
| Propanoate metabolism | X | X | X | Facilitation |
| *Lipid metabolism* | | | | |
| Fatty acid metabolism | X | Facilitation | X | X |
| Fatty acid degradation | Facilitation | X | Facilitation (n.s.) | Facilitation |
| Fatty acid biosynthesis | Facilitation | X | X | X |
| Lipid biosynthesis proteins | X | Facilitation | X | X |
| *Glycan biosynthesis and metabolism* | | | | |
| Lipopolysaccharide biosynthesis | X | X | Selection (n.s.) | X |

Table 2: **Summary of KEGG pathways (ii)** showing significant differences between phases in their mean proportions (rows) for the three datasets considered (columns). The stages compared in each dataset are indicated. Cells indicate the phase in which the mean proportion is significantly higher, having an "X" if no significant differences were found. Cells labelled as (n.s.) indicate that the difference is not significance when corrected for multiple testing. This table summarizes pathways classified in KEGG level 1 as Metabolism. Carbohydrate and lipid metabolism are most represented in the facilitation phase, while nitrogen metabolism, metabolism of cofactor and vitamins, glycan biosynthesis and nucleotide metabolism are relatively higher at the selection or attachment phases.

| | Metagenomes | | Isolates | |
|---|---|---|---|---|
| | Experiments | PiCRUST | | |
| Pathway / Stages compared | Selection / facilitation | Selection / facilitation | Selection / facilitation | Attachment / facilitation |
| **Metabolism** | | | | |
| *Amino acid metabolism* | | | | |
| Arginine biosynthesis | Selection | X | X | X |
| Arginine and proline metabolism | X | X | X | Facilitation |
| Valine, leucine and isoleucine degradation | Facilitation | Facilitation | Facilitation (n.s.) | Facilitation |
| Lysine degradation | Facilitation | X | Facilitation | Facilitation |
| Glycine, serine and threonine metabolism | X | X | Facilitation (n.s.) | X |
| Phenylalanine metabolism | X | X | X | Facilitation |
| beta-Alanine metabolism | X | X | X | Facilitation |
| Tryptophan metabolism | Facilitation | Facilitation | Facilitation | Facilitation |
| Histidine metabolism | X | X | Facilitation (n.s.) | X |
| *Xenobiotics metabolism* | | | | |
| Aminobenzoate degradation | X | Facilitation | X | X |
| Benzoate degradation | X | X | Facilitation | Facilitation |
| *Unclassified* | | | | |
| Other metabolic pathways | X | Facilitation | X | X |

Table 3: **Summary of KEGG pathways (iii)** showing significant differences between phases in their mean proportions (rows) for the three datasets considered (columns). The stages compared in each dataset are indicated. Cells indicate the phase in which the mean proportion is significantly higher, having an "X" if no significant differences were found. Cells labelled as (n.s.) indicate that the difference is not significance when corrected for multiple testing. This table summarizes pathways classified in KEGG level 1 "Metabolism". Amino acid and xenobiotic metabolism are more represented in bacteria at the facilitation phase.

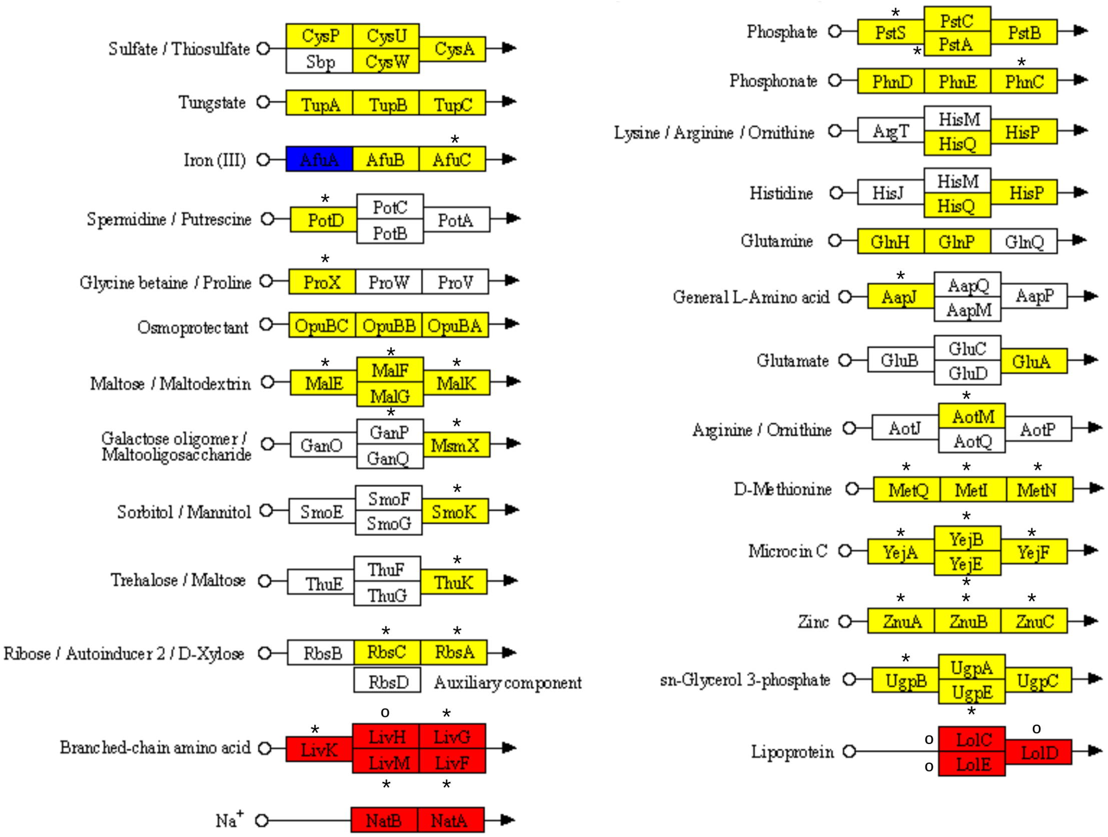

Figure 13: **ABC transporters.** The nodes highlighted contain genes whose proportion is significantly higher in the metagenome predictions for samples classified at the attachment (yellow), or facilitation stages (red). An asterisk (∗) near a gene's box indicate that the isolates with a preference for the stage indicated in the color of the box have in their genome the gene, and that the proportion of isolates having the gene is larger than in groups of isolates with other preferences, i.e. both metagenome predictions and genome content in the isolates are consistent. If this group of isolates have the gene but their proportion is not the largest, a circle (∘) is displayed.

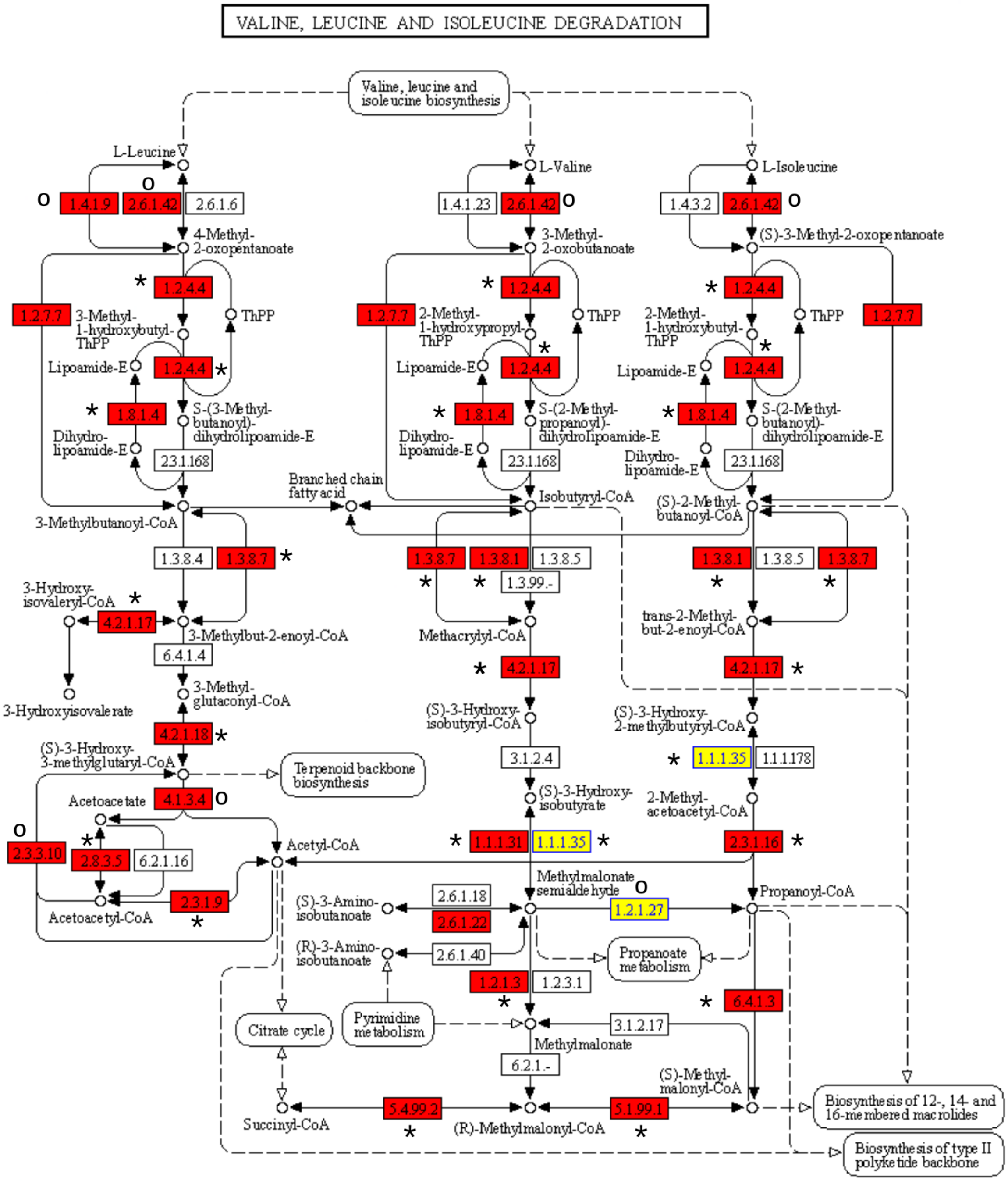

Figure 14: **Valine, leucine and isoleucine degradation.** The nodes highlighted contain genes whose proportion is significantly higher in the metagenome predictions for samples classified at the attachment (yellow), or facilitation stages (red). An asterisk (∗) near a gene's box indicate that the isolates with a preference for the stage indicated in the color of the box have in their genome the gene, and that the proportion of isolates having the gene is larger than in groups of isolates with other preferences, i.e. both metagenome predictions and genome content in the isolates are consistent. If this group of isolates have the gene but their proportion is not the largest, a circle (○) is displayed.

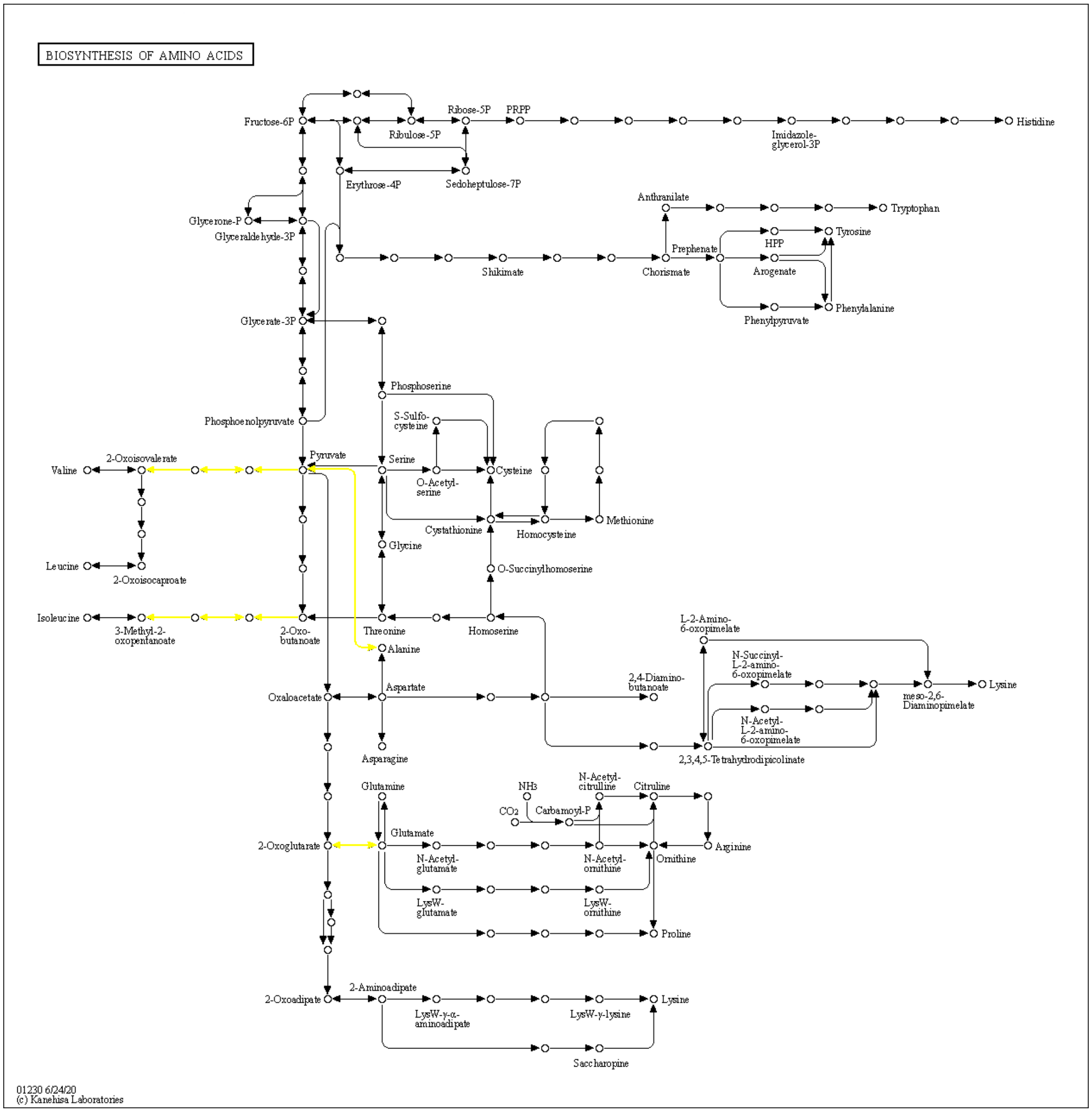

Figure 15: **Valine, leucine and isoleucine biosynthesis.** The pathways highlighted contain genes whose proportion is significantly higher in the metagenome predictions for samples classified at the attachment stage (yellow).

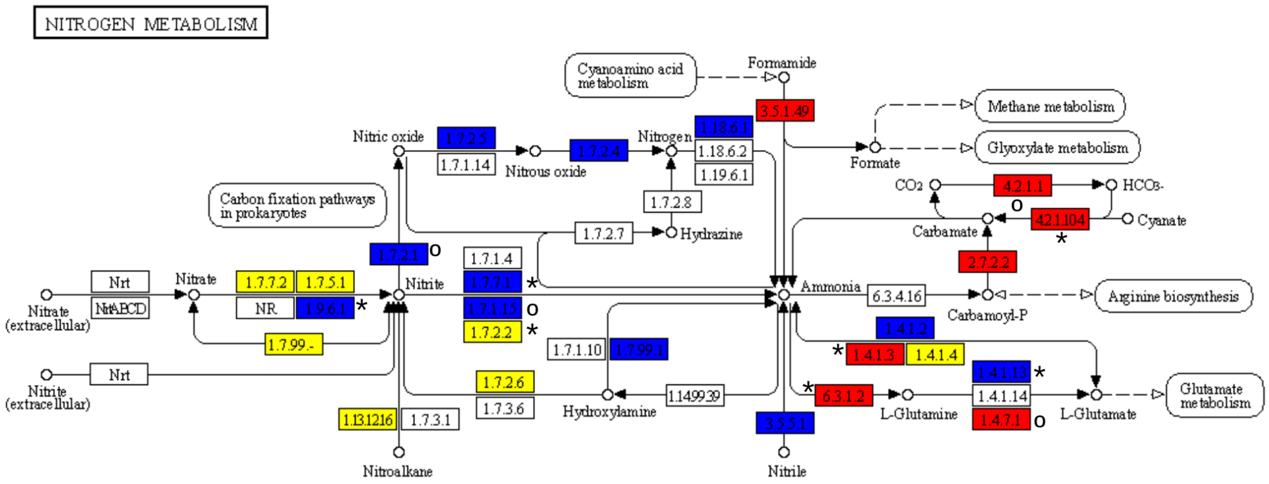

Figure 16: **Nitrogen metabolism.** The nodes highlighted contain genes whose proportion is significantly higher in the metagenome predictions for samples classified at the attachment (yellow), selection (blue) or facilitation stages (red). An asterisk (∗) near a gene's box indicate that the isolates with a preference for the stage indicated in the color of the box have in their genome the gene, and that the proportion of isolates having the gene is larger than in groups of isolates with other preferences, i.e. both metagenome predictions and genome content in the isolates are consistent. If this group of isolates have the gene but their proportion is not the largest, a circle (∘) is displayed.

#### 2 Supplementary Note: Validation of PiCRUST predictions

The analysis of metagenome sequencing experiments revealed that several samples had a low number of reads which, since we sequenced only one replicate per substrate and time-point, reduced our chances to find significant differences between sets of samples. To complement this information we performed a prediction from the 16S rRNA amplicon sequences with PiCRUST [2], from which three replicates per sample and time-point were available. This allowed us to obtain better statistics for any comparison between sets of samples. We computed the quality of PiCRUST predictions quantifying the NSTI score [2] from which we retrieved a median of 0.089 across samples, a value close to the lower bound found for human samples, and at the upper bound of mammal samples (see Fig. 3 in Ref. [2]).

We further compared the experimental metagenomic samples having a number of annotated genes in KEGG $>$5K with its correspondent prediction. 75% of the genes found in the experiments were predicted, with a median of the Spearman's correlation coefficient between the experimental and predicted samples of 0.57, suggesting a fair agreement. Next we investigated in more detail if the biological picture provided was comparable, analysing which are the differences between samples obtained at early times and late times for the experimentally-measured metagenomes, finding 17 pathways with significant differences that we took as reference (presented in the Main Text). Proceeding similarly with PiCRUST predictions we found that 8 pathways were represented in the predictions with similar quantitative values and always consistently predicting an enrichment towards the same time-window (Main Text).

We then explored the remaining 9 pathways to understand the discrepancies. We analysed each pathway individually now considering the attachment, selection and facilitation stages. For each pathway, we performed an individual ANOVA test, followed by a Bejamini-Hochber-corrected Tukey-Kramer post-hoc test, and filtering pathways with $\eta > 0.2$. We found 5 more pathways consistent with the experimental metagenomics (e.g. "Mismatch Repair", see Suppl Fig. 17) and 1 more that predicted significantly enriched genes in the attachment phase instead of in the selection phase (ABC transporters, Suppl. Fig. 17, note that the attachment phase was not considered in the experimental metagenomes). Therefore only 3 pathways from experimental data showing significant differences between early and late stages were not found significant in the predictions (arginine biosynthesis, porphyrin/clorophyll metabolism and aminoacyl t-RNA biosynthesis) and we did not find any pathway with a prediction contradicting the experimental data.

The validated pathways were considered the core pathways to build the main picture discussed in the Main Text. In addition, PiCRUST predictions brought a much larger number of pathways with statistically significant differences between early and late time-windows (although without verification these could be false positives). 114 pathways were identified after removing pathways belonging to the categories "human diseases" and "organismal systems" (e.g. bacterial invasion of epithelial cells). Finally, some of these significant pathways that were consistent with the broad picture built with the experimentally-validated pathways and having high effect sizes (e.g. nitrogen metabolism $\eta = 0.47$) were incorporated in the results discussed in the Main Text.

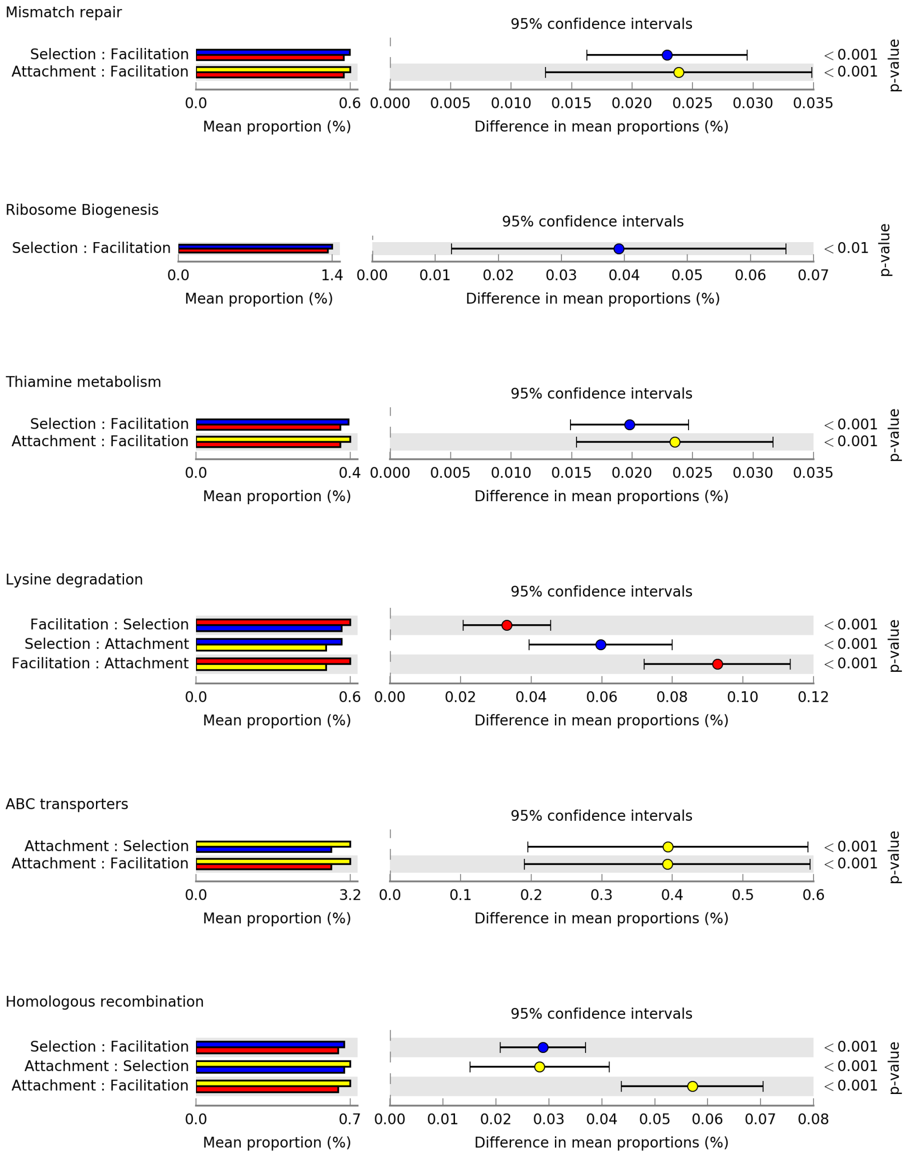

Figure 17: **Post-Hoc tests of individual pathways**. The analysis test significant differences in the mean proportion of the number of genes predicted with PiCRUST, for samples belonging to two out of the four time-windows identified : "attachment" stage (yellow), "selection" or "early" stage (blue) and "facilitation" or "late" stage (red). Each row represents a pairwise test between two of these time-windows. Only significant tests out of the 6 possible tests per pathway ae shown (Benjamini-Hochberg corrected Tukey-Kramer test, $p < 0.05$). The seven pathways were found to have significant differences between early and late stages in the experimental metagenomics. All pathways are consistent except ABC transporters, that have most significant differences with respect to the "attachment" stage, which is not considered in the metagenomics analysis and we analyse in more detail in the Main Text.

#### References

[1] Pascual-García A, Bonhoeffer S, Bell T. Metabolically cohesive microbial consortia and ecosystem functioning. Philosophical Transactions of the Royal Society B. 2020;375(1798):20190245.

[2] Langille MG, Zaneveld J, Caporaso JG, McDonald D, Knights D, Reyes JA, et al. Predictive functional profiling of microbial communities using 16S rRNA marker gene sequences. Nature Biotechnology. 2013;31(9):814–821.
